# Extracellular communication between brain cells through functional transfer of Cre mRNA

**DOI:** 10.1101/2023.01.29.525937

**Authors:** David Rufino-Ramos, Kevin Leandro, Pedro R.L. Perdigão, Killian O’Brien, Maria Manuel Pinto, Magda M. Santana, Thomas S van Solinge, Shadi Mahjoum, Xandra O Breakefield, Koen Breyne, Luís Pereira de Almeida

**Author notes:** Corresponding authors: Luís Pereira de Almeida, Ph.D., CNC—Center for Neuroscience and Cell Biology, University of Coimbra, Rua Larga, Coimbra 3004-504, Portugal, Koen Breyne, Ph.D., Molecular Neurogenetics Unit, Massachusetts General Hospital, 149 Thirteenth Street, Charlestown, MA 02129 USA, Correspondence and requests for materials should be addressed to K.B. or to L.P.d.A. These authors contributed equally to this work (David Rufino-Ramos, Kevin Leandro). These authors contributed equally to this work (Xandra O Breakefield, Koen Breyne, Luís Pereira de Almeida).

## Abstract

In the central nervous system (CNS), the crosstalk between neural cells is mediated by extracellular mechanisms, including brain-derived extracellular vesicles (bdEVs).

To study endogenous communication across the brain and periphery, we explored Cre-mediated DNA recombination to permanently record the functional uptake of bdEVs cargo overtime. To elucidate functional cargo transfer within the brain at physiological levels, we promoted the continuous secretion of physiological levels of neural bdEVs containing Cre mRNA from a localized region in the brain by *in situ* lentiviral transduction of the striatum of Flox-tdTomato Ai9 mice reporter of Cre activity. Our approach efficiently detected in vivo transfer of functional events mediated by physiological levels of endogenous bdEVs throughout the brain. Remarkably, a spatial gradient of persistent tdTomato expression was observed along the whole brain exhibiting an increment of more than 10-fold over 4 months. Moreover, bdEVs containing Cre mRNA were detected in the bloodstream and extracted from brain tissue to further confirm their functional delivery of Cre mRNA in a novel and highly sensitive Nanoluc reporter system.

Overall, we report a sensitive method to track bdEVs transfer at physiological levels which will shed light on the role of bdEVs in neural communication within the brain and beyond.

## 2 INTRODUCTION

Exchange of nucleic acids, proteins and lipids between brain cells, including neurons, oligodendrocytes, astrocytes and microglia are an essential aspect of homeostasis in the brain (Q. Li and Barres 2018). Cell-to-cell communication in the CNS is crucial to support the function and integrity of neurons, control inflammation and mediate the removal of debris and infectious agents (Chu and Williams 2021)(Hill 2019). Intercellular transfer of molecules in the brain is mediated through direct cytoplasmic connections between cells, such as tunneling nanotubes (TNTs) (Khattar et al. 2022; Rustom et al. 2004), and paracrine communication mediated by the release of signaling molecules, such as growth factors, neurotransmitters and cargo of extracellular vesicles (EVs) (Hill 2019). EVs are nano-sized particles naturally produced by all cells and surrounded by a lipid bilayer, which protects their contents from degradation (van Niel, D’Angelo, and Raposo 2018).

EVs are virtually released by all neural cells to neighboring or distant compartments and influence a wide range of processes throughout the body (Hill 2019)(Shi et al. 2019). In fact, EVs released by different cell types of the nervous system have been shown to have different cell binding specificities and fates for their cargo upon internalization. For example, while neuroblastoma-derived EVs were predominantly internalized by glia, those secreted by cortical neurons were preferentially taken up by other neurons (Chivet et al. 2014). On the other hand, oligodendrocyte EVs were shown to be internalized by neurons and microglia and contribute to neuronal integrity (Frühbeis et al. 2013). In fact, essential roles for neural function and integrity have been attributed to EV communication between neurons and other neural cells, including synapse assembly and plasticity, neuronal survival and immune responses (Budnik, Ruiz-Cañada, and Wendler 2016)(Zappulli et al. 2016). In addition, EVs have been shown to be involved in several neurodegenerative diseases, such as Alzheimer’s and Parkinson’s disease (Emmanouilidou et al. 2010; Hill 2019; Mahjoum, Rufino-ramos, and Broekman 2021; Rajendran et al. 2006; Wang et al. 2017).

EVs communicate between neural cells by transferring protein and nucleic acids cargo to recipient cells which alters their gene expression and function. Among nucleic acids, small DNA fragments (Balaj et al. 2011)(Liu et al. 2022) and different extracellular RNA (exRNAs) species have been found, such as mRNAs, microRNA, piwi-interacting RNAs and other non-coding RNAs ((Nolte-’t Hoen et al. 2012; Wei et al. 2017). In fact, there are reports supporting transfer of mRNAs <3 kb with some efficiency. Nevertheless, enrichment of long RNA sequences is reduced due to packaging limitations (O’Brien et al. 2020). RNA transfer within the brain has been involved in regulation of gene expression in astrocytes (Morel et al. 2013), decreased (Bhaskaran et al. 2019; Lang et al. 2012) or accelerated (Abels, Maas, et al. 2019; Van Der Vos et al. 2016) glioma growth and spreading of misfolded proteins in Alzheimer’s and Parkinson’s disease (Wang et al. 2017)(Rajendran et al. 2006)(Emmanouilidou et al. 2010).

Thus, uncovering the roles of EVs and their functional events in the CNS is crucial to better understanding of their function in neuronal physiology and pathological conditions. To obtain insights into brain communication mediated by EVs in disease and non-disease conditions, approaches based on extracting bdEVs from brain tissue have been used to obtain insights into brain communication mediated by EVs and their composition. Despite some reports suggesting L1CAM and NCAM as promising candidates to isolate bdEVs, there is no consensus yet about a specific neuronal marker to selectively isolate bdEVs in the bloodstream (Norman et al. 2021; Ter-Ovanesyan et al. 2021). Interestingly, lipidomics (Su et al. 2021), proteomics (Huang et al. 2020; Vassileff et al. 2020)(You et al. 2022) and transcriptomics (Huang et al. 2020) were performed to investigate the profile of bdEVs signatures to further discriminate normal and diseased EVs produced in neurodegenerative diseases. Despite the findings of some dysregulated molecules with potential to serve as brain signatures, there is a current need to understand whether the bdEVs directly isolated from brain tissue truly represent the EVs population secreted from cells.

Studying biological functions of EVs *in vivo* present limitations regarding the need of large amount of previously concentrated particles exposed to cells (Gupta, Zickler, and El Andaloussi 2021). Despite showing functional transfer of proteins and exRNAs that result in phenotypic changes on recipient cells (Pegtel et al. 2010; Valadi et al. 2007), there is a lack of understanding the physiological role of EVs transfer. In fact, studying functional activity of EV carrying exRNAs at physiological levels in the brain is challenging for several reasons, such as the low number of RNA molecules per vesicle (Albanese et al. 2021; M. Li et al. 2014; Wei et al. 2017), the degradation of RNA transferred cargo which hinders the identification of EV-mediated effects in recipient cells, and the lack of definitive specific markers to isolate bdEVs (You et al. 2022).

To counteract these limitations, targeting genomic DNA using systems such as Cre-LoxP reporters induces permanent changes at the DNA level allowing a permanent recording of functional events mediated by rare endogenous EV (Van Duyne 2015), and offers the advantage of validation with multiple readout analyses from DNA to RNA and protein levels. Fluorescent reporter genes, such as tdTomato are typically used under the control of promoters with stop regions between LoxP sites that are removed upon Cre activation (Madisen et al. 2010). Fluorescent genes can be replaced by luminescent reporters which possess high sensitivity, such as Nanoluciferase (Nanoluc)(Hall et al. 2012)(England, Ehlerding, and Cai 2017).

Cre-LoxP based systems are powerful systems to study EV-mediated intercellular cargo transfer both *in vitro* and *in vivo* (Frühbeis et al. 2013; Ridder et al. 2015; Sterzenbach et al. 2017)(Zomer et al. 2015). Cre mRNA was shown to be naturally incorporated into EVs without requirement of packaging signals (Steenbeek et al. 2018; Zomer et al. 2015). Functional transfer of Cre molecules contained in EVs was shown to be essential in discriminating metastatic behavior *in vivo* (Ruivo et al. 2022; Zomer et al. 2015) by marking cells which internalized vesicles through the expression of fluorescent proteins (Zomer et al. 2016), suggesting the possibility of applying the same rational to study brain communication.

In this study, we aimed at studying brain communication mediated by endogenous bdEVs secreted from the striatum to peripheral brain regions. We generated an *in vivo* brain region continuously secreting bdEVs carrying Cre mRNA upon intracranial injection of lentiviral vectors (LVs) encoding the Cre transgene into the striatum of Ai9 reporter mice. Upon transduction, striatal cells continuously express and package Cre molecules in bdEVs. The continuous exposure of brain cells to bdEVs containing Cre mRNA resulted in an increase of tdTomato signal in the whole mouse brain from 4 weeks to 16 weeks as a consequence of a spatial gradient from the initial injection site of LVs and the continuous spreading of bdEVs carrying Cre mRNA over time. Through this strategy we demonstrated EV-mediated brain communication by permanently recording at the DNA level the continuous uptake of their functional cargo in the brain. Moreover, we showed that bdEVs can be isolated from brain tissue samples or the bloodstream and successfully internalize into neurons *in vitro* to functionally deliver Cre mRNA.

## 3 RESULTS

### 1 Extracellular communication is shown through the functional transfer of Cre activity *in vitro*

In this study, we aimed at studying brain communication mediated by EVs. For that purpose, we developed a reporter system based on the Cre-LoxP recombination which allows detection and recording of rare events mediated by extracellular communication through genomic DNA recombination. Therefore, HEK293T cells were used as a continuous source of EVs packaging Cre mRNA after stable transduction with a lentiviral vector encoding CRE sequence, under the control of a phosphoglycerate kinase (PGK) promoter. A firefly luciferase (Fluc) reporter under the Ubiquitin C gene (UbC) promoter was included as an indicator for Cre expression. Both promotors are ubiquitously expressed and ensured stable and high levels of expression in EV donor cells (Wettergren et al. 2012)(Norrman et al. 2010), generating applicability to a wide variety of cell types. To retain the protein products of both transgenes in the donor cells, a nuclear localization signal (NLS) and a H2B histone was added to the N-terminal of the CRE and Fluc genes (Figure 1A). Cre (Figure 1A) and Fluc (Supplementary Figure 1A) protein content were mainly restricted to the nucleus of transduced HEK293T cells. Fluc expression resulted in over 5-fold increase in bioluminescence of transduced HEK293T cells but was barely detectable in culture media (Supplementary Figure 1B).

**Figure 1.**
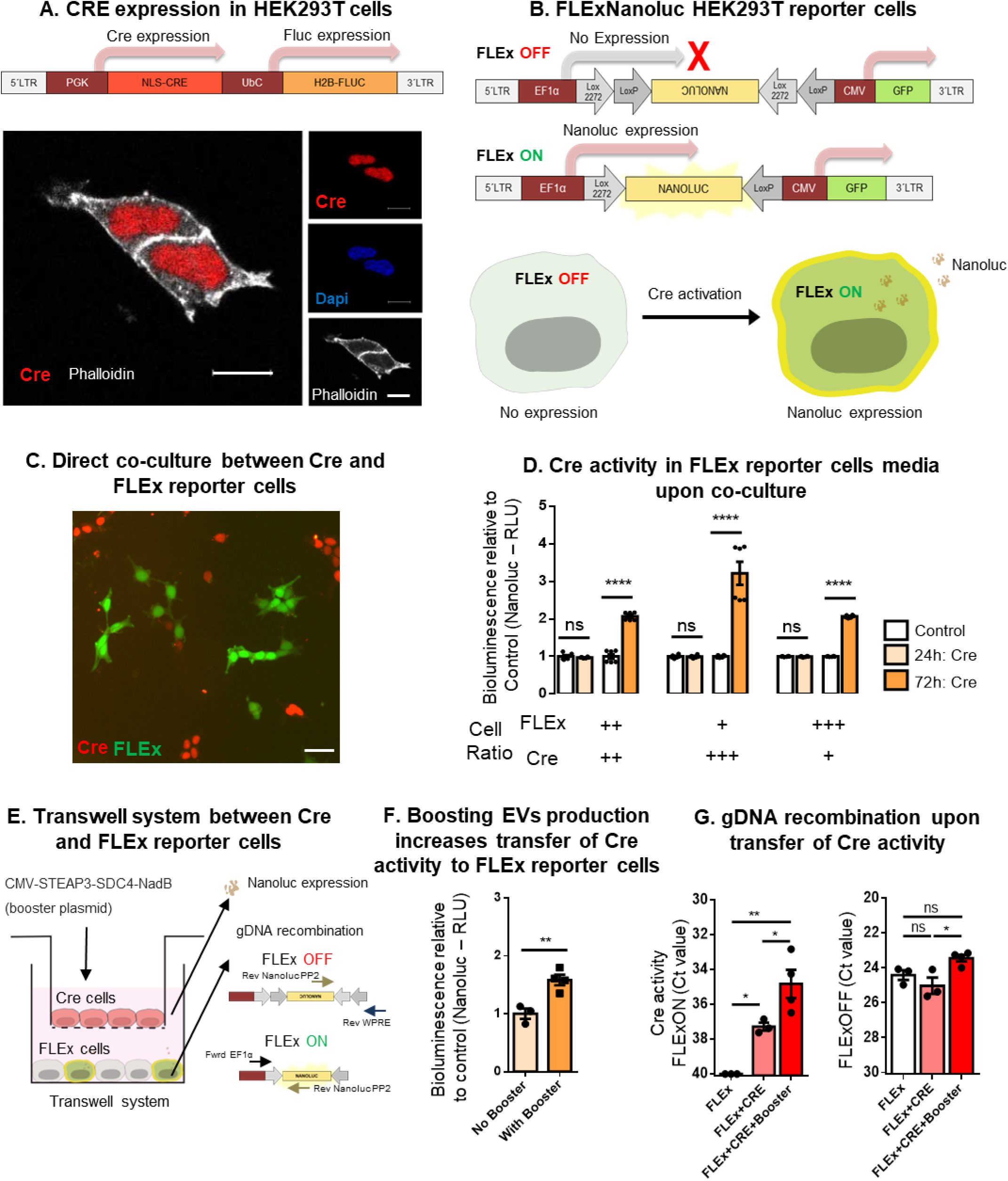
Extracellular communication shown through functional transfer of Cre activity *in vitro*. A. (Top) Schematic representation of the lentiviral construct expressing NLS CRE (1026bp) under control of PGK promoter, and H2B firefly luciferase (Fluc) (1650bp) under control of UBC promoter. Cre and Fluc genes contain a nuclear localization signal (NLS) and H2B, respectively, at the N-terminus that shuttles the proteins to the nucleus. (Bottom) Representative immunofluorescent image from confocal microscopy of HEK293T cells stably expressing Cre protein (red) mainly in the nucleus (blue). Actin filaments in cytoplasm were stained with Phalloidin (white). Scale bar, 10 µm. B. Schematic representation of FLExNanoluc switch used to generate a sensitive Cre reporter system. The FLExNanoluc in the OFF-state does not allow Nanoluciferase (Nanoluc) expression, because the gene is backwards in the construct. Upon Cre activation the Nanoluc gene flips and becomes in frame with the EF1α promoter in the ON-state. The resulting Nanoluc expression generates detectable bioluminescence in both cells and media. C. Co-culture of HEK293T cells stable expressing Cre (red) and HEK 293T cells stable expressing FLExNanoluc and GFP (green) for 72h. Scale bar represents 20 µm. D. Bioluminescence evaluation of Nanoluc secreted in media. Nanoluc signal in the cell media detected after 24 and 72 hours of co-culture. Cells were cultured in three FLExNanoluc:Cre ratios (1:1; 1:3 and 3:1). The white bars represent a control condition in which FLExNanoluc reporter cells were co-cultured with WT HEK293T cells (no expression of Cre). Cre activity is represented by bioluminescence signal relative to control (N=6). Data is presented as mean ± SEM and compared by Unpaired t test, ****p < 0.0001. E. Transwell system (1μm pore inserts) with Cre cells seeded on the apical side of the upper chamber and previously transfected with CMV-STEAP3-SDC4-NadB plasmid to boost small EV production and FLExNanoluc reporter cells seeded in the lower chamber, with the latter showing recombination mediated by EVs. F. Cre activity in boosted condition relative to non-boosted condition is represented by Nanoluc bioluminescence (RLU) in FLEx cells (N=3). Data presented as mean ± SEM and compared by Unpaired t test **p < 0.01. G. Evaluation of gDNA recombination by RT-PCR showing Ct values of non-recombined DNA (FLExOFF) and recombined DNA (FLExON) (N=3/4). FLEx condition (white bar) was used to establish a baseline condition corresponding to no recombination. Data represented as Ct values obtained in each sample condition. Data is presented as mean ± SEM and compared by one-way ANOVA followed by Tukey’s multiple comparison test (F = 19.72, F = 6.956), *p < 0.05 and **p < 0.01.

To detect the EV-mediated extracellular transfer of Cre and permanently register its activity in the genomic DNA (gDNA), we generated a reporter cell line based on the FLExNanoluc switch reporter, (Breyne et al. 2022). In the OFF-state, the FLExNanoluc system does not express Nanoluciferase (Nanoluc) as its coding sequence is flipped between LoxP regions (Figure 1B). In the presence of Cre, the Nanoluc gene is flipped to the ON-state following the inversion and excision of the flanking LoxP sites, thus allowing the Nanoluc gene to be expressed by restoring its frame with the upstream human elongation factor-1 alpha (EF1 α) promoter (Figure 1B). A constitutively active CMV-driven GFP reporter was placed downstream of the floxed region to monitor cells encoding the FLEXNanoluc system. Following co-transfection with FLEXNanoluc and Cre-expressing constructs, the resulting Nanoluc expression generates detectable bioluminescence signal upon addition of furimazine substrate, both in cells (Supplementary Figure 1C) and media (Supplementary Figure 1D). FLEx reporter activation increased proportionally with the amount of Cre activity encoded by 10ng, 50ng and 100ng PGK-Cre-UbC-Fluc plasmid (Supplementary Figure 1D). HEK293T cells were transduced with a lentiviral vector encoding EF1α FLExNanoluc to generate stable-expressing reporter cell lines. To evaluate whether Cre exRNA was transferred from the donor cell line to the FLExNanoluc recipient cells, we co-cultured both cell types in three FLEx:Cre ratios, as represented in Figure 1C. Immunocytochemistry allowed distinction of donor and recipient cells using anti-Cre antibody staining (red) and GFP endogenous expression from the FLEx construct (Figure 1C). In the control condition, non-transduced HEK293T cells were co-cultured with FLEx reporter cells. After 24 hours of co-culture, bioluminescence was not significantly different from controls. In contrast, after 72 hours of co-culture with transduced HEK293T cells, a significant increase in Nanoluc bioluminescence was observed in all conditions, with a 2-fold increase between the 1:1 and 3:1 FLEx/Cre ratios, and more than 3-fold increase for 1:3 FLEx/Cre ratio (Figure 1D). Data suggests that Cre activity is dependent on time and dose to mediate Nanoluc activation.

To exclude the possibility that the observed effects could result from direct cell fusion or formation of tunnelling nanotubes (TNTs), we used a transwell system permeable only to particles less than the 1μm pore size to restrict exchange between donor and recipient cells to EVs. Cre exRNA donor cells were seeded on the apical side of the upper chamber and FLEx recipient cells were seeded in the lower chamber. (Figure 1E). To boost EV production, we transfected donor cells with the CMV-STEAP3-SDC4-NadB plasmid (Kojima et al. 2018), a tricistronic expression construct described as regulating three distinct pathways - exosome biogenesis, budding of endosomal membranes to form multivesicular bodies and cellular metabolism - which overall increases the production of small EVs (Kojima et al. 2018). Cre activity mediated by EVs between boosted and non-boosted Cre cells was evaluated by bioluminescence of the FLExNanoluc reporter cells. Boosted Cre cells presented 1.58-fold increase in bioluminescence as compared to non-boosted condition (Figure 1F). To validate Cre-mediated FLEXNanoluc activation, gDNA recombination was analyzed using a RT-qPCR strategy that allows distinguish between recombined and non-recombined DNA. The levels of non-recombined DNA (FLExOFF) were similar between conditions (roughly Ct values of 24). In contrast, the levels of recombined DNA (FLExON) were found to be significantly higher in the condition where Cre donor cells were boosted for production of EVs (Figure 1G). Overall, these results suggest that extracellular communication was mediated through functional transfer of Cre species in particles sized below 1μm, presumably extracellular vesicles.

### 2 Cre activity is mediated by transfer of Cre mRNA through EVs, but not Cre protein

In order to investigate whether EVs transfer Cre molecules, we isolated EVs from culture media of Cre expressing cells after 72 hours by Size Exclusion Chromatography (SEC) (Benedikter et al. 2017). After pelleting cell debris and concentrating media, we resolved the samples in SEC columns and collected EV fractions 7 to 11 (Figure 2A), as previously described (Théry et al. 2018). EVs were analyzed for their protein content by western blotting and found to be positive for typical protein EV markers, including Alix, HSC70 and TSG101 and negative for the endoplasmic reticulum marker, calnexin and Cre protein (Figure 2B). The latter was detected in Cre expressing cells but not in their derived EVs, possibly due to the predominant localization of Cre in the cell nucleus due to the presence of the NLS (Figure 1A). Importantly, Cre mRNA was detected both in donor cells (Ct value of 22) (Supplementary Figure 2A) and in their derived EVs (Ct value of 25) (Figure 2C), being packaged in these vesicles. Similarly, Fluc mRNA was detected in cells (Supplementary Figure 2A) and their derived EVs (Supplementary Figure 2B).

**Figure 2.**
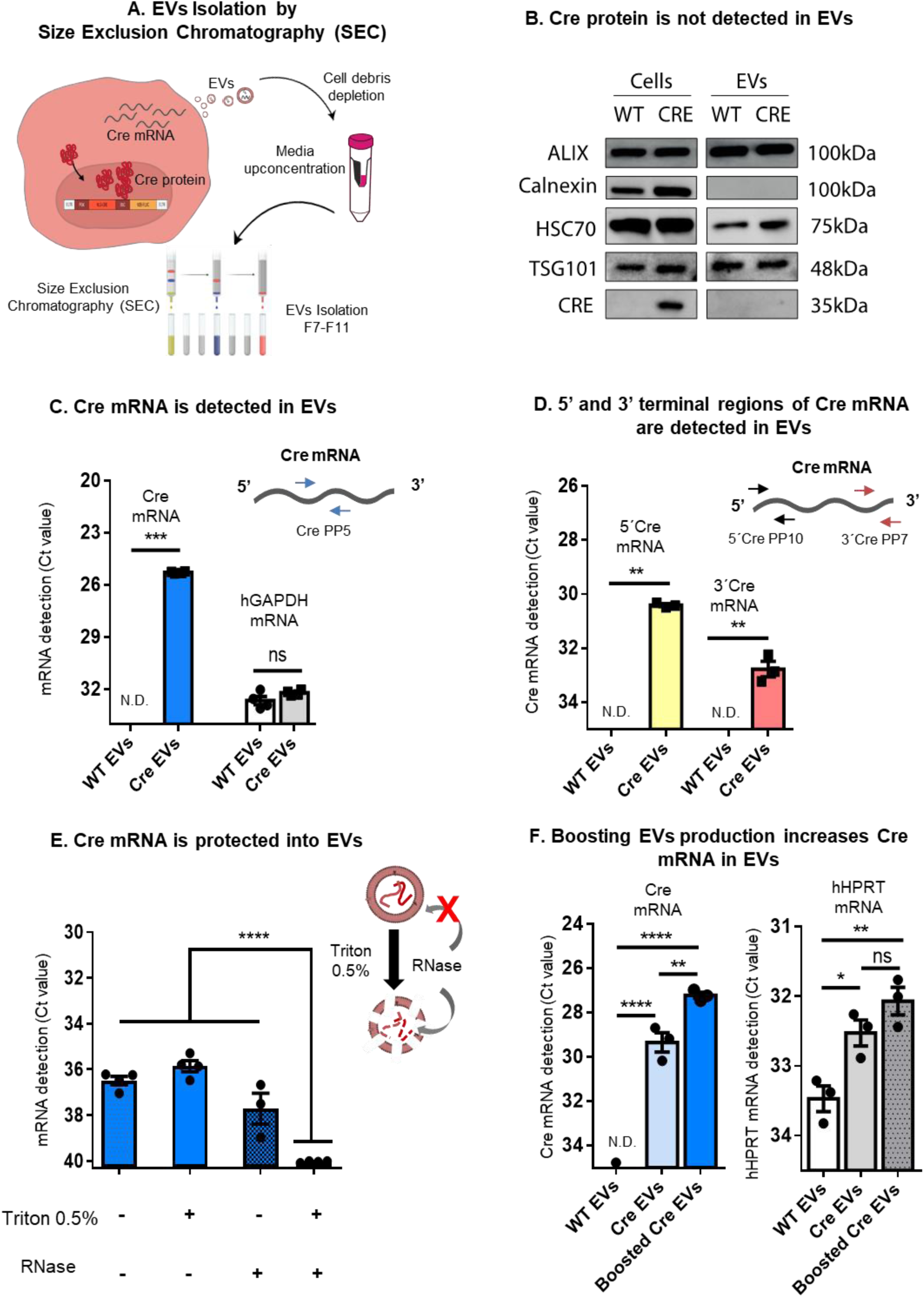
Cre activity is mediated by transfer of Cre mRNA through EVs. A. Schematic representation of EVs isolation by Size Exclusion Chromatography (SEC). Briefly, EVs were isolated from the media of HEK293T stably expressing Cre, cell debris were removed (300gx10minutes) and media concentrated (100kDa filter) to a final volume of 500uL and then loaded onto a qEV Original SEC column. 5 EV-enriched fractions of 500 µL were collected (fractions 7-11). B. Western blotting of equimolar amounts of protein from cells and their derived EVs shows the positive markers Alix, HSC70 and TSG101 and undetectable levels of the ER marker calnexin. Cre protein is present in Cre donor cells but was not detectable in EVs from those cells. C. Cre mRNA is detected in Cre EVs, but not WT EVs (N=4). hGAPDH was detected in both conditions. Data presented as Ct values - mean ± SEM and compared by Unpaired t test with Welch’s correction. Statistical significance: ***p < 0.001 and ns – nonsignificant. D. 5’ and 3’ regions of Cre exRNA are detected in Cre EVs, but not in WT EVs (N=3). Data is presented as Ct values mean ± SEM and compared by Unpaired t test with Welch’s correction. Statistical significance: **p < 0.01 E. Cre EVs treated with RNase A in the presence or absence of 0.5% Triton X-100 showed Cre-exRNA is predominantly protected inside EVs (N=4). Data is presented as mean ± SEM and compared by ordinary one-way ANOVA followed by Dunnett’s multiple comparison test (F=493.4). Statistical significance: ****p < 0.0001. F. CMV-STEAP3-SDC4-NadB booster plasmid increases EV production and Cre exRNA detection. hHPRT was used as a housekeeping control. Data is presented as mean ± SEM and compared by ordinary one-way ANOVA followed by Sidak’s multiple comparisons test (F=192.4). Statistical significance: *p < 0.05, **p < 0.01, ***p < 0.001 and ns – nonsignificant.

To further investigate to what extent Cre mRNA is loaded into EVs we used different Cre primer pairs (PP) targeting the 5’ or 3’ ends of Cre mRNA. We detected a higher Ct value in the 5’ region (Ct value of 30) when compared to the 3’ region (Ct value of 32) (Figure 2D), suggesting a mixture of Cre mRNAs were present in EVs, and possibly Cre mRNA is degraded starting from 3’ region (Houseley and Tollervey 2009). To evaluate whether Cre mRNA is protected within the EV lumen, EVs were exposed to 0.5% Triton X-100 and RNase, which disrupts EV membranes and degrades mRNA, respectively. Treatment with either RNAse or 0.5% Triton alone did not significantly reduce Cre mRNA Ct value. In contrast, Cre mRNA was not detected when EVs were exposed to both RNAse and 0.5% Triton (Figure 2E), which supports Cre mRNA being protected in the lumen of EVs.

The overexpression of CMV-STEAP3-SDC4-NadB plasmid in cells boosted small EVs production, in terms of particle numbers and CD63 species (Supplementary Figure 2C). To evaluate whether Cre mRNA was packaged into small EVs, Cre-expressing cells were transfected with CMV-STEAP3-SDC4-NadB or control plasmids (CMV-RFP), EVs isolated and Cre mRNA analyzed. Boosting EVs production with CMV-STEAP3-SDC4-NadB plasmid increased the detection of Cre mRNA in EVs to a Ct value of 27, when compared to non-boosted Cre EVs (transfected with a control plasmid of CMV-RFP) for which a Ct value of 29 was found, suggesting small EVs originating from the endocytic pathway contain Cre mRNA. The lower Ct value of Cre mRNA in boosted EVs relates to a higher secretion of EVs from boosted cells as observed from higher particle count (Supplementary Figure 2C), as compared to Ct values of HPRT in boosted and non-boosted EVs (Figure 2F).

Overall, our data indicates that we have established a Cre expressing cell line continuously secreting EVs which have a natural ability to package Cre mRNA but not NLS-modified Cre protein.

### 3 EVs transfer functional Cre mRNA *in vitro* and *in vivo*

To investigate whether Cre mRNA detected in EVs would be functionally transferred to recipient cells, we exposed FLExNanoluc reporter cells to EVs isolated from Cre mRNA donor cells for 24 and 72 hours. To determine Cre activity, Nanoluc bioluminescence was evaluated in culture medium 24 and 72 hours after incubation (Figure 3A). The first 24h of incubation led to a 10% increase while the 72h of incubation led to a 50% increase in bioluminescence relative to control (incubation with HEK293T EVs), suggesting a time-dependent effect.

**Figure 3.**
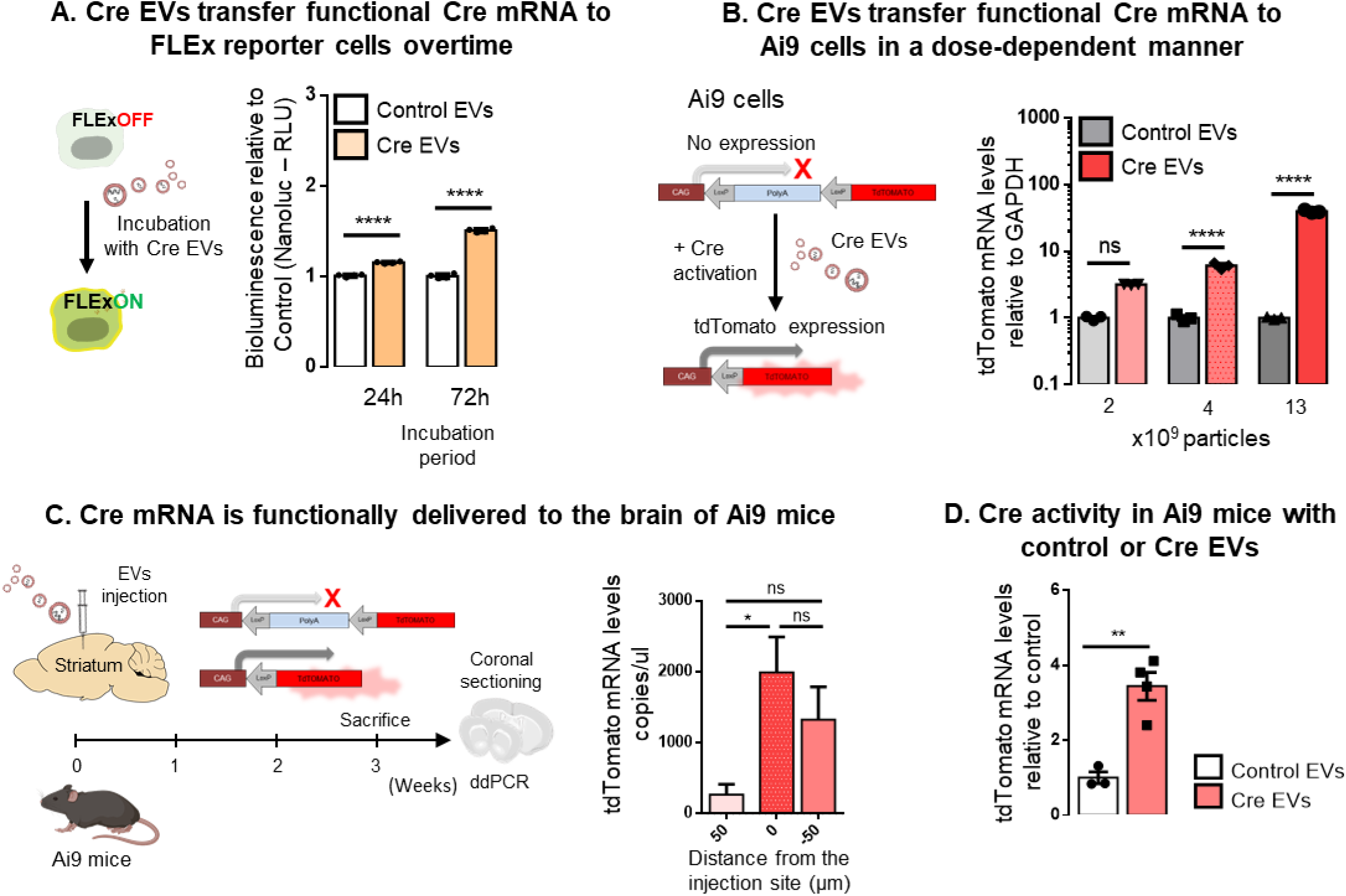
Concentrated EVs transfer functional Cre mRNA *in vitro* and *in vivo*. A. Cre EVs transfer functional Cre mRNA to FLEx reporter cells overtime. FLEx reporter cells were incubated with Cre EVs and Nanoluc bioluminescence evaluated in culture medium 24 and 72 hours after incubation. Cre activity is represented by bioluminescence signal relative to control (incubated with WT EVs). Data presented as means ± SEM and compared by Unpaired t test. Statistical significance: ****p < 0.0001. B. Cre EVs transfer functional Cre exRNA to Ai9 cells in a dose-dependent manner. Schematic illustration of Ai9 reporter in which tdTomato expression is prevented by a stop cassette between the promoter and the coding sequence. Removal of the stop cassette by Cre activation results in tdTomato expression. Bar graphs represent tdTomato expression levels evaluated by RT-digital droplet PCR (ddPCR) post-incubation with three different doses of Cre-EVs (2.2, 4.4 and 13.1 X10^9^ particles) for 72 hours. Data presented as means ± SEM and compared Unpaired t test. Statistical significance: ****p < 0.0001. C. Cre mRNA is functionally delivered to the brain of Ai9 mice. Schematic illustration of Cre EVs intracranially injected in Ai9 reporter mice. Three weeks post-injection, tdTomato mRNA levels in coronal brain sections were evaluated through ddPCR to detect the injection site of Cre EVs (N=4). Data is presented as tdTomato copies/uL mean ± SEM and compared by one-way ANOVA followed by Tukey’s multiple comparisons test (F=5.641). Statistical significance: *p < 0.05 and ns – nonsignificant. D. Cre activity of exogenous EVs in brain. Control EVs (from HEK293T) or Cre EVs injected intracranially into Ai9 mice were compared for Cre activity in the coronal sections at the injection site in the brain. tdTomato expression at the injection site in the striatum of animals were evaluated by Droplet Digital PCR (ddPCR) (Control N=3 and Cre EVs N=4). Data is presented as mean ± SEM and compared by Unpaired t test. Statistical significance: **p < 0.01.

To evaluate whether Cre activity induced by Cre EVs uptake is dependent on the dose, we used the Ai9 reporter cells in which tdTomato expression is prevented by a stop cassette between two LoxP sites (floxed), encoding three tandem polyA sequences between the chicken β-actin (CAG) promoter and the gene coding sequence (Madisen et al. 2010). The removal of the stop cassette upon Cre activation results in tdTomato expression. Co-transfection of HEK293T cells with Ai9 plasmid and increasing amounts of Cre plasmids led to an increase in tdTomato mRNA levels, also validated by detection of gDNA recombination (Supplementary Figure 3A). To evaluate functional Cre mRNA transfer through EVs, Ai9 reporter cells were incubated with three different doses of Cre EVs (2.2, 4.4 and 13.1 X10^9^ particles) for 72 hours and mRNA expression evaluated by digital droplet PCR (ddPCR). The lowest dose of EVs (2.2×10^9^ particles) resulted in 3.1 times significantly higher tdTomato expression than the control (incubation with HEK293T EVs), while the intermediate dose (4.4×10^9^ particles) resulted in 6 times higher tdTomato expression and the highest dose (13.1×10^9^ particles) resulted in 39 times higher tdTomato expression (Figure 3B). These results were confirmed at the DNA level (Supplementary Figure 3B). Together, these results indicate a dose dependent effect of Cre EVs on tdTomato signal *in vitro*.

To investigate whether Cre EVs mediate functional transfer of Cre mRNA *in vivo*, we injected 1×10^9^ particles of Cre EVs intracranially into the striata of Ai9 transgenic mice (Figure 3C). The same number of HEK293T EVs (lacking Cre mRNA) were injected in the same region of Ai9 mice. Three weeks after the injection, mice were sacrificed and ddPCR of striatum coronal sections was performed to determine the injection. For that aim, we monitored the levels of tdTomato expression in each coronal region that corresponds to the peak of Cre activity in the brain (Figure 3C). To evaluate if the increase in tdTomato signal was due to functional delivery of Cre mRNA, we compared the tdTomato mRNA levels in brain sections of animals injected with Cre EVs and control EVs (Figure 3D). TdTomato mRNA levels of animals injected with Cre EVs were 3.5-fold higher (50µm sections) relative to control animals, suggesting tdTomato expression at the injection site was dependent on the activity of Cre mRNA functionally delivered by EVs. Overall, our *in vitro* and *in vivo* data indicates that EVs conveying Cre mRNA are responsible for Cre-mediated activity detected by the LoxP reporter *in vitro* and *in vivo*.

### 4 Neurons establish a long-term source of Cre mRNA within the brain

To unravel the role of the spreading EVs in brain communication *in vivo*, we generated a brain endogenous Cre secreting region in the striatum using LVs encoding PGK-driven Cre and UbC-driven Fluc genes (Figure 4A). Fluc bioluminescence was used to monitor gene expression in transduced brain cells in living mice (Figure 4B). A bioluminescent signal was observed upon intraperitoneal (I.P.) injection of D-luciferin (100mg/kg) in LV Cre-injected mice, but not in control mice (injected with 1% PBS/BSA). Bioluminescence was used to monitor gene expression overtime (1, 2, 3, 4, 8, 12 and 16 weeks) without the need to sacrifice the mice. When compared to the control injected animals, there was a tendency for increased Fluc bioluminescence signal over time, suggesting that donor cells are not removed from the brain after transduction and are thus a stable source of Cre expressing cells to further produce bdEVs able to stably secrete endogenous EVs containing Cre (Figure 4B).

**Figure 4.**
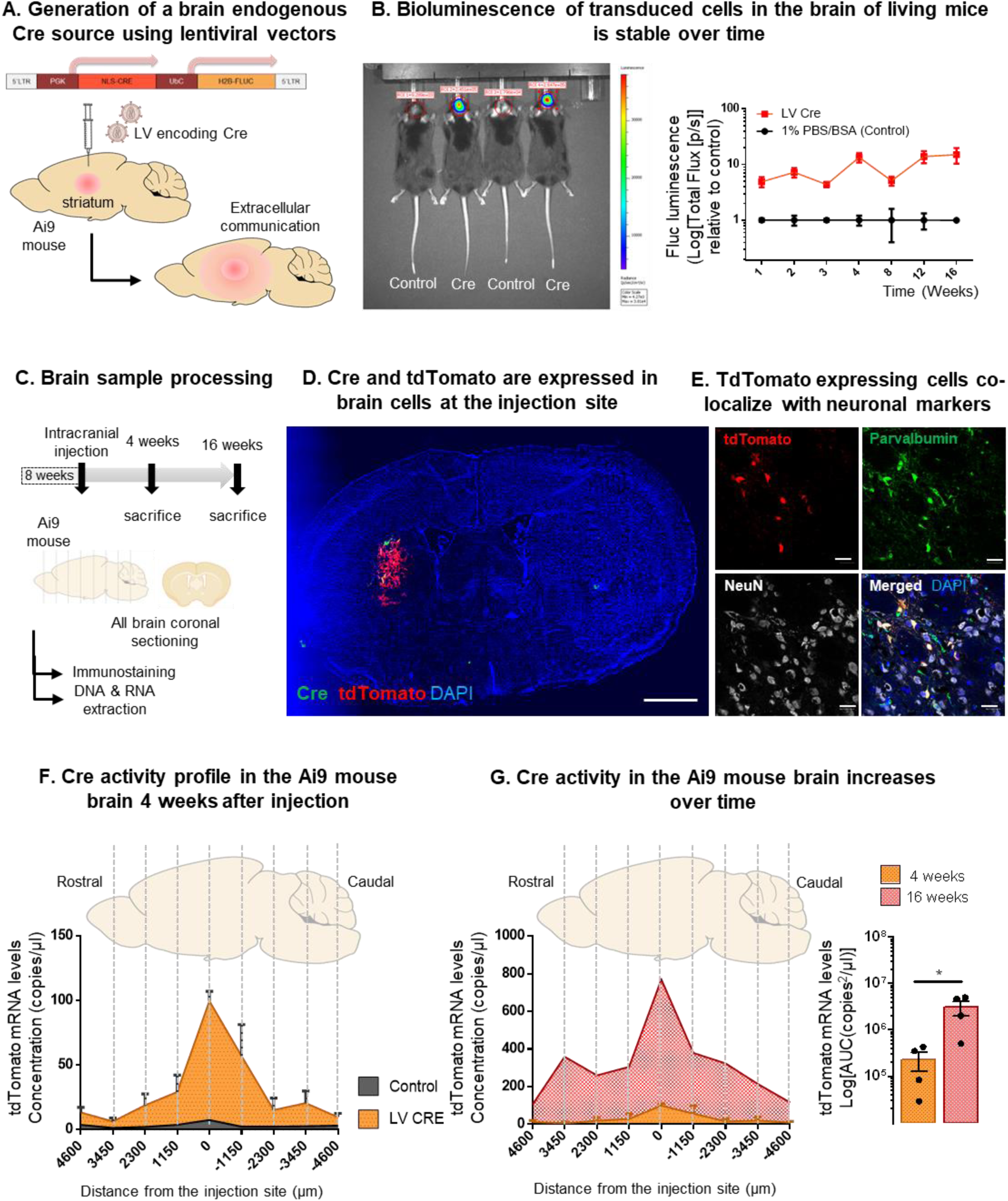
Cre activity within the brain is shown through long term transduction of neurons *in vivo*. A. Generation of an endogenous brain source of Cre EVs upon intracranial injection of lentiviral vectors (LVs) into the striatum of Ai9 mice. B. Firefly luciferase bioluminescence was used to monitor transduced brain cells in living mice. Stable production of Cre and Fluc in the brain was monitored by bioluminescence *in vivo* from 1 to 16 weeks following intracranial injection of LVs. C. Brain sample processing. Ai9 animals intracranially injected with LV encoding Cre were sacrifice 4-and 16-weeks post injection. Whole brain coronal sectioning was performed, and sections processed for immunostaining or DNA/RNA extraction. D. Immunofluorescence of coronal sections at the injection site at 4 weeks post-intracranial transduction. Brain cells expressing Cre (green) and tdTomato (red) upon intracranial injection of lentivirus encoding Cre in the striatum. Analysis performed with a Keyence BZ-X810 microscope 20x (injection site, scale bar 200μm). E. tdTomato positive cells co-localize with parvalbumin and NeuN suggesting the majority of the transduced cells are inhibitory neurons. Nucleus is represented by DAPI staining. Images are representative of a group of five Ai9 animals. Analysis performed with a laser confocal microscopy equipped with Plan-Apochromat 40×/1.40 Oil DIC M27 (420782-9900) (neurons, scale bar 20μm). F. Cre activity profile in the Ai9 mouse brain 4 weeks after LV injection. Whole-brain coronal sections were used to compare tdTomato mRNA expression levels in the brains of Ai9 mice injected with LV Cre (orange) or 1%PBS/BSA (grey). The highest tdTomato signal was detected at the injection site, decreasing in distal rostral and caudal regions (N=4). Data presented as tdTomato copies/ul means ± SEM. G. Cre activity in the Ai9 mouse brain increases over time. Comparison between tdTomato expression in the whole brain of LV Cre injected mice after 4 weeks (orange) or 16 weeks (red). Area under the curve (AUC) of tdTomato expression among the two conditions is shown in copies x µm/µl means ± SEM and compared by Unpaired t test. Statistical significance: *p < 0.05.

To evaluate the extent of EV communication throughout the brain, Ai9 animals were injected intracranially with LVs encoding Cre and sacrificed at 4- and 16-weeks post injection. Whole brain coronal sectioning was performed from the rostral to caudal regions and sections processed for immunostaining or DNA/RNA extraction (Figure 4C). The Cre source in the brain was characterized by immunofluorescence on coronal sections in the striatum of Ai9 animals 4 weeks after intracranial injection (Figure 4D). Cre expressing cells were found to co-localize with tdTomato expressing cells resulting from Ai9 reporter activity (Supplementary Figure 4A). TdTomato-positive cells expressed the neuronal makers NeuN and parvalbumin (Figure 4E) and MAP2 (Supplementary Figure 4B), suggesting that neurons also expressed PGK-Cre-UbC-Fluc. Moreover, GFAP-positive cells were found to partially co-localize with tdTomato positive cells, suggesting astrocytes were also partially transduced (Supplementary Figure 4C). In contrast, IBA1-positive cells did not co-localize with tdTomato expressing cells (Supplementary Figure 5C), suggesting microglia was not transduced by viral vectors or it migrated to the injection site after the viral injection activity period.

To evaluate whether Cre activity diffuses from the injection site through extracellular mechanisms, longitudinal tdTomato expression profiles from the whole brain were investigated. To that purpose, coronal sections of Ai9 mice with a total thickness of ∼160 µm were collected from the rostral to the caudal region of the brain to extract RNA for ddPCR analysis. To evaluate whether extracellular mechanisms have a significant impact on tdTomato expression throughout the brain over time, we compared tdTomato expression in different sections from the whole brain 4 and 16 weeks after injection. At 4 weeks, we observed a high level of tdTomato at the injection site (0 µm) with a 14-fold increase in expression between control and LV-Cre injected mice with from 7 to 100 copies mRNA/µL (Figure 4F), while at adjacent (distance 1150 µm) and peripheric regions (distance >3450 µm) tdTomato levels were restricted to <75 and <20 copies/µL, respectively.

Remarkably, at 16 weeks post-injection, we observed a 10-fold increase of tdTomato expression across all brain sections compared to 4 weeks (Figure 4G), with an area under the curve of 2, 749,457 copiesxµm/µL (16 weeks) compared with 268,460 copiesxµm/uL (4 weeks). The highest tdTomato expression was still observed at the injection site, nonetheless we observe a significant increase in the reporter levels at 16 weeks in the adjacent and peripheric sections compared to 4 weeks post-injection. These results suggest an increase in the spatial gradient of Cre activity overtime, possibly due to extracellular mechanisms including EVs diffusion and transfer of Cre mRNA to peripheral brain regions. Moreover, It is unlikely to be mediated by the contribution of LVs spreading from the injection site, since they are not capable of replication and their half-life in culture is less than 48 hours according to (Dautzenberg, Rabelink, and Hoeben 2021). Although tdTomato expression is primarily driven by Cre activation of LoxP sites, we cannot exclude the spreading of secondary bdEVs carrying tdTomato molecules from floxed cells. To exclude the contribution of bdEVs spreading reporter species, we physically separated two distinct brain regions: striatal donor cells secreting bdEVs carrying Cre mRNA and cerebellar recipient cells containing the reporter system, even though we were not successful in showing functional transfer of Cre activity over this extensive distance (Supplementary Figure 5D, 5E and 5F).

More interestingly, we observed that bdEV communication is not restricted to the brain, as we were able to detect diffusion of EVs produced in the brain compartment into the bloodstream. SEC was used to isolate EVs from serum, followed by immunoprecipitation of EVs containing CD9, CD63 and CD81 (Supplementary Figure 5B and 5C). Cre mRNA was detected in immunoprecipitated SEC EVs from serum of mice injected in the brain with LVs Cre as compared to their injected controls, corroborating our hypothesis that brain EVs can transfer Cre mRNA from the injection site to other regions and confirming the extension of bdEV communication beyond the brain.

Overall, these results suggest that localized sustained *in vivo* neuronal secretion of EVs induces effects in extended brain regions that accumulate over time.

### 5 Brain tissue derived-EVs (bdEVs) deliver functional Cre mRNA

To provide further evidence that extracellular transfer of Cre molecules from the striatum were mediated to some extent by bdEVs, we next set out to isolate these bdEVs by adapting previously described protocols (Huang et al. 2020; Su et al. 2021, 2021; Vella et al. 2017). For this purpose, we digested brains of LV Cre-injected mice with collagenase type III and isolated bdEVs according to their density and size, as described in Figure 5A. BdEVs were isolated by Optiprep^TM^ (Iodixanol) Density Gradient (ODG, Supplementary Figure 6A) and 10 fractions were collected according to their densities (Figure 5B) - fraction 1 corresponds to the lower density fraction (1.02 g/mL) and fraction 10 to the higher density fraction (1.25 g/mL). Then, we evaluated the protein amount in each fraction (Figure 5C). Fraction 1 corresponded to 32% of total protein decreasing to 1.8% in fraction 10, suggested that protein distribution in fractions decreased as density increased. After applying a downstream ultracentrifugation step (100,000 g for 2 hours) of each individual fraction, the protein profile changed favoring bdEVs isolation (Figure 5C). This latter purification step eliminated free protein contamination in the first fractions as demonstrated by our micro bicinchoninic acid (microBCA) measurements before and after ultracentrifugation (Figure 5C). The higher percentage of protein was found in ODG fractions 6, 7 and 8 (25%, 12% and 14%) from the bdEV associated pellet at 100,000g (Théry et al. 2018), with reduction of free proteins in the first fractions (fraction 1, 2 and 3 – 5%, 9%, 13%, respectively). These results emphasize the relevance of ultracentrifugation as a final step to wash and concentrate EVs. This protocol was used in the subsequent experiments to isolate bdEVs.

**Figure 5.**
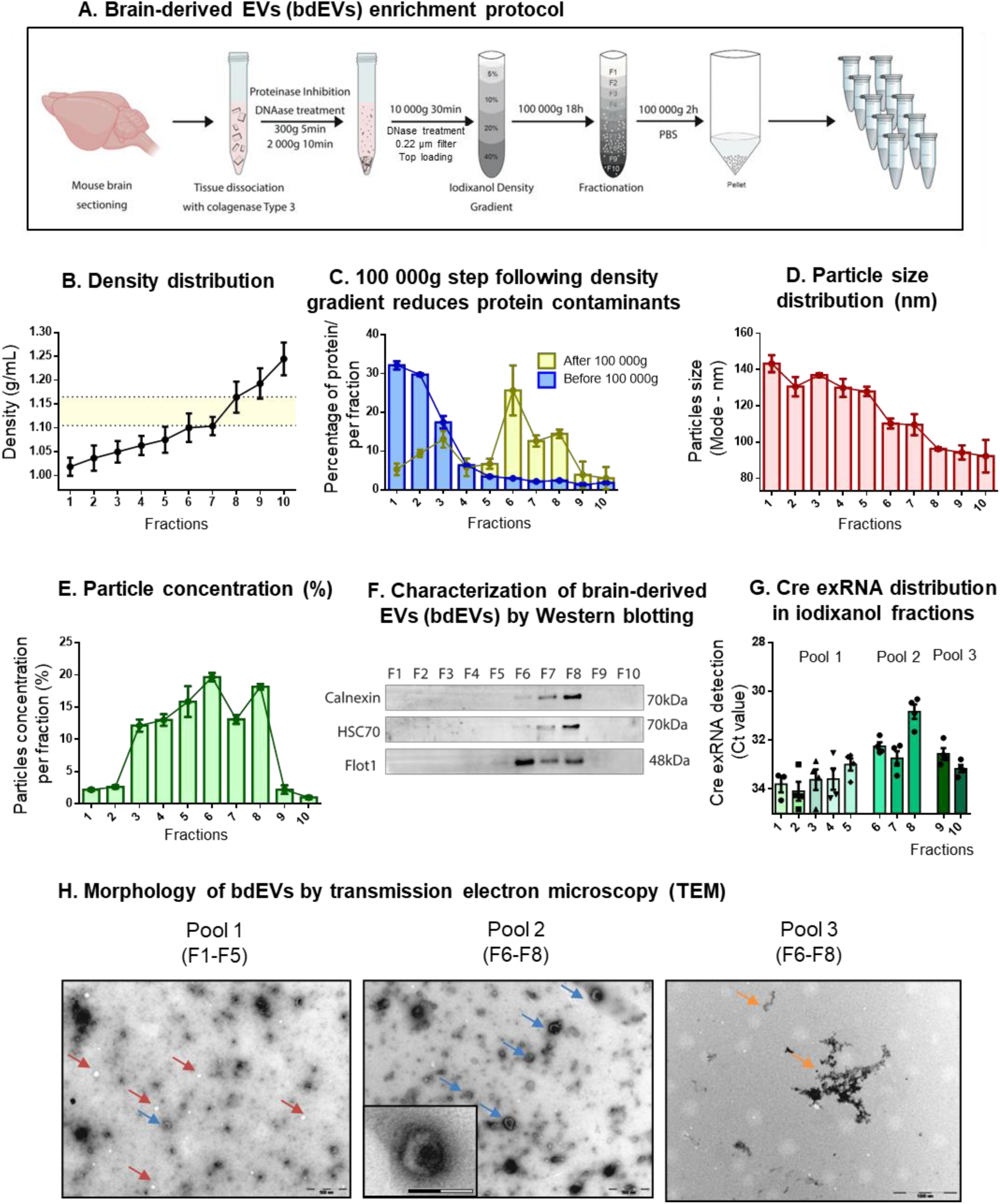
Cre mRNA is detected in brain derived-EVs (bdEVs) extracted from the brain. A. Schematic illustration of the protocol used to isolate brain-derived EVs (bdEVs). B. Density distribution of 10 fractions as result of iodixanol gradient centrifugation at 100,000g for 18 h. EV-enriched fractions were isolated in densities ranging from 1.105 to 1.165 g/mL (midle region) (N=10). C. Quantification of protein amount per fraction (in percentage) before and after 100,000g purification step. Before 100,000g purification step (blue bars), protein is highly enriched in the first fractions decreasing until fraction 10. After 100,000g purification step (yellow bars), the majority of free protein was washed out and the highest percentage of protein was located in EV fractions 6,7 and 8 (N=4). D. Particle size distribution of each fraction (represented by mode) was evaluated by Nanoparticle tracking analysis (NTA) (red bars). Fraction 1 showed the higher mode with 140nm and decreasing in each fraction until fraction 10 that showed the mode of 90nm (N=3). E. Particle concentration in each fraction was evaluated by NTA (green bars), with fractions 6, 7 and 8 accounting for more than 50% of total particles, while fractions 1 and 2, and 9 and 10 showed a lower concentration (N=3). F. Representative western blotting of 10 fractions obtained after ODG and ultracentrifugation of each fraction in PBS (loaded per volume) show the presence of positive EV markers HSC70 and flotilin-1. The endoplasmic reticulum protein calnexin was detected in low levels in EV-enriched fractions. G. Distribution of Cre exRNA in bdEV fractions was evaluated by RT-qPCR (Ct Value). Fractions 6, 7 and 8 showed higher levels of Cre exRNA when compared to the other fractions (N=4) (same volume was used as starting point). H. Transmission electron microscopy (TEM) of pool 1 (fraction 1-5) showed lipoproteins (red arrow) and few canonial bdEVs (blue arrow), pool 2 (fraction 6-8) was highly enriched in bdEVs (blue arrow) with cup-shaped format, and pool 3 (fraction 9-10) presented very low number of particles and some protein aggregates (orange arrows). Scale bars are 500nm (big pictures) and 200nm (Pool 2, Crop). Values are presented as mean ± SEM.

BdEVs were characterized in terms of particle size mode (Figure 5D) and particle concentration (Figure 5E) assessed by Nanoparticle Tracking Analysis (NTA), we observed that particles from fraction 1 presented the higher particle size mode (140nm), which gradually decrease until fraction 10 with the lowest size mode (90nm) (Figure 5D and Supplementary Figure 6B). Particle concentration was higher in middle fractions, particularly in fraction 6 and fraction 8 which represented 19.6% and 18.1% of the total particle concentration, respectively (Figure 5E). Interestingly, fractions 6, 7 and 8 accounted for more than 50% of total particles, which was increased by the 100,000g ultracentrifugation step.

We corroborated bdEVs isolation through this method by western blotting for total protein (Supplementary Figure 6C) and specific EV protein markers (Figure 5F). Fractions 6, 7 and 8 were positive for HSC70 (70kDa) and Flotilin-1 (48kDa). Interestingly, the protein calnexin (70kDa) was detected in low levels in the EV-enriched fractions, suggesting this type of EVs are made at contact sites with the endoplasmic reticulum as suggested in (Barman et al. 2022). Following the confirmation that EVs were derived from brain tissue, we aimed to evaluate the distribution of Cre exRNA in all fractions by RT-PCR. Interestingly, Cre exRNA was detected in EV enriched fractions 6 (Ct=32), 7 (Ct=33) and 8 (Ct=31), as compared to other fractions (Figure 5G). A similar profile was detected when Fluc mRNA was analyzed in bdEVs (Supplementary Figure 6D).

Taking these findings into consideration, we grouped the 10 fractions in 3 different pools based of particles characteristics (Figure 6A). We analyzed the size and concentration profile of each pool by NTA (Supplementary Figure 6B) and performed transmission electron microscopy (TEM) to access EVs morphology (Figure 5H). Pool 1 (fractions 1-5) was highly enriched in lipoproteins (red arrows), showing few canonical EVs, pool 2 (fractions 6-8) was highly enriched in EVs with cup-shaped morphology and pool 3 (fractions 9-10) was depleted of EVs, showing mostly protein aggregates.

**Figure 6.**
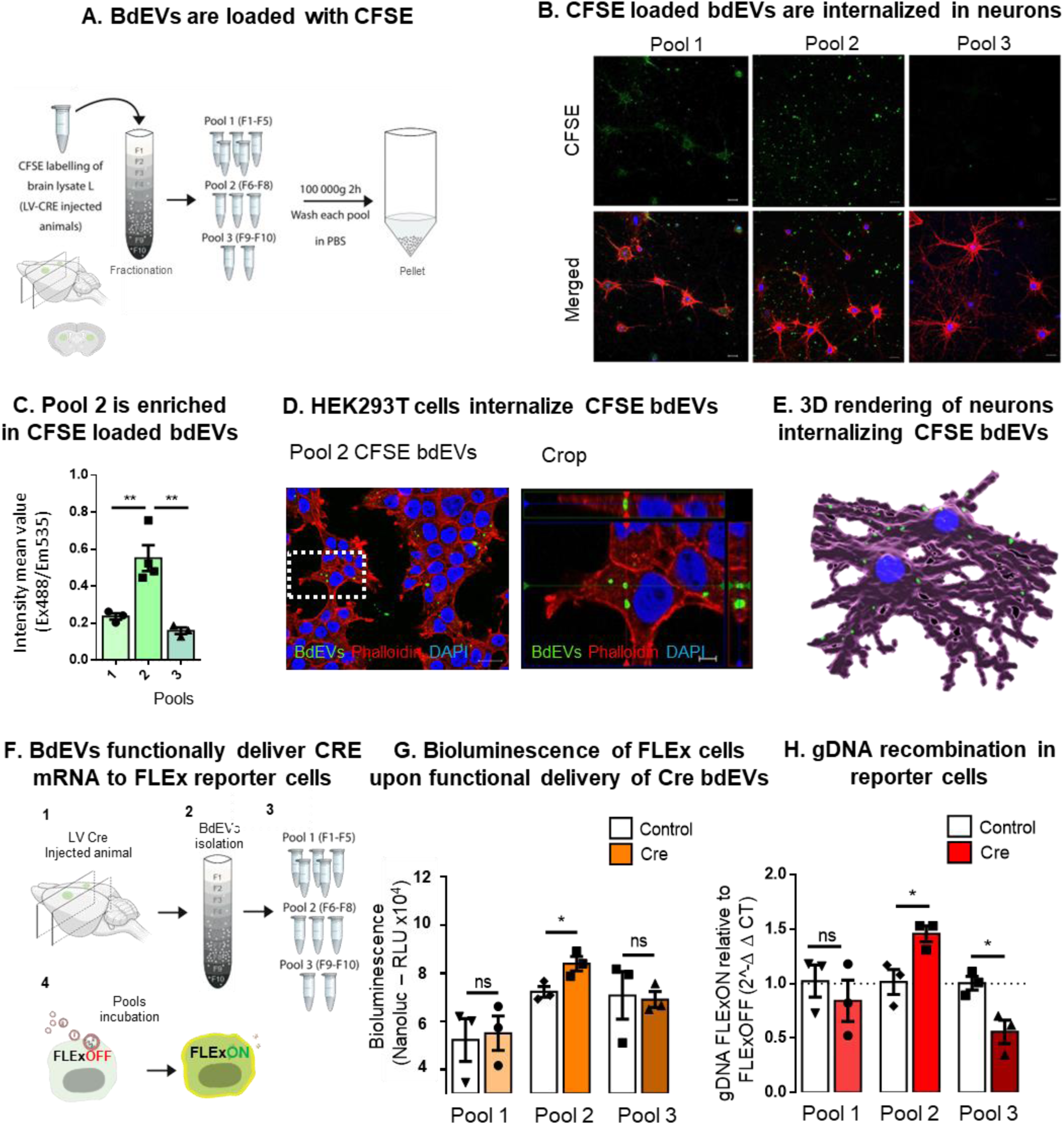
Brain derived-EVs (bdEVs) are taken up by neurons and deliver functional Cre mRNA. A. Schematic illustration of the protocol used to isolate bdEVs labelled with carboxyfluorescein succinimidyl ester (CFSE) from Cre injected mice. Thick coronal sections containing the injection sites were used as starting material for the EV extraction. B. CFSE loaded bdEVs were exposed to neurons. The 10 fractions of CFSE labelled EVs were divided in 3 pools: pool 1 (fraction 1-5), pool 2 (fraction 6-8) and pool 3 (fraction 9-10) after density gradient separation. Each pool was incubated with cultured primary hippocampal neurons and total CFSE fluorescence was measured. Scale bar 5 μm. C. Pool 2 presented the highest fluorescence signal when compared to the other two pools (N=3/4). Data presented as means ± SEM and compared by ordinary one-way ANOVA followed by Dunnett’s multiple comparisons test (F=17.41). Statistical significance: **p < 0.01. Scale bar 20 μm. D. Incubation of Pool 2 of CFSE labelled bdEVs (green) with HEK293T cells (red) in culture (left), followed by high magnification image (right) of primary neurons internalizing bdEVs (green). Cells were stained with phalloidin (red) and DAPI (blue) and analyzed by laser confocal microscopy equipped with Plan-Apochromat 40×/1.40 Oil DIC M27 (420782-9900). Scale bars - 20 μm (left) and crop (right) 5 μm. E. Imaris 3D rendering showing individual brain-derived EVs (green) being internalized in primary hippocampal neurons in culture (Scale bar 20 μm). F. Schematic representation of bdEVs delivering functional Cre mRNA to FLExNanoluc reporter cells. G. Detection of Cre activity by measurement of Nanoluc bioluminescence in FLExNanoluc reporter cells. The same number of particles was incubated in control (white bars) and Cre conditions (orange bars). The highest luminescent peak was detected in pool 2 containing Cre when compared to control pool 2 carrying the same number of bdEVs without Cre. Values are presented as mean ± SEM. Unpaired t test was used to evaluate statistical significance: * p ≤ 0.05 and ns for non-significant. H. Detection of Cre activity was confirmed at DNA level by analyzing the ratio between FLExON (recombined) and FLExOFF (non-recombined) between Control and Cre samples. Values are presented as mean ± SEM. Using unpaired t test, statistical significance: * p ≤ 0.05 and ns for non-significant.

Considering the successful isolation of bdEVs from brain tissue, our next goal was to evaluate whether retrieved vesicles were functional and could be effectively taken up by recipient brain cells. BdEVs were labelled with a green fluorescent dye, carboxyfluorescein succinimidyl ester (20 µM CFSE) and isolated by ODG followed by ultracentrifugation (Figure 6A). Upon measuring CFSE fluorescence in all fractions, we observed a gradual decrease in fluorescence from fraction 1 to fraction 7, with an increase in fraction 8 (Supplementary Figure 6E). We hypothesized that fluorescence from the first fractions corresponded to free CFSE molecules nonspecifically bound to lipoproteins, while the peak in fraction 8 corresponds to CFSE incorporated into bdEVs (Supplementary Figure 6E).

To evaluate whether CFSE-labelled bdEVs could be internalized by recipient cells, the three pools described above: pool 1 (fractions 1-5), pool 2 (fractions 6-8) and pool 3 (fractions 9-10) were incubated with HEK293T cells (Supplementary Figure 6H) and primary hippocampal rat neurons (Figure 6B). The uptake was assessed by measuring the fluorescence intensity after 6 hours of incubation with bdEVs. Pool 2, corresponding to bdEVs, presented over 2-fold increase in fluorescent signal with a mean value of 0.55 a.u., when compared to pool 1 with mean value of 0.23 a.u. and pool 3 with a mean value of 0.16 a.u (Figure 6C, \*\**p* < 0.01), suggesting cells exposed to pool 2 took up CFSE-labelled particles. Additionally, confocal microscopy confirmed CFSE-labelled bdEVs internalization by recipient HEK293T cells (Supplementary Figure 6H) and primary neurons (Supplementary Figure 6G) after 6 hours of incubation. Moreover, a high magnification image showed that bdEVs accumulated in the cytoplasm of HEK293T cells (Figure 6D). Similar results were observed in rat primary neuronal cultures, where CFSE-labelled bdEVs (Pool 2) were efficiently internalized (Supplementary Figure 6G). A 3D rendering reconstitution of primary hippocampal neurons internalizing bdEVs from pool 2 showed individual bdEVs inside cells, particularly present in neuronal extensions (Figure 6E).

To study bdEV fate post-uptake, pools of bdEVs isolated from brains of Cre and control injected mice were isolated, as described above and incubated with FLExNanoLuc reporter cells (Figure 6F). Following 72h, cells were analyzed for Nanoluc bioluminescence as result of Cre activity. We found no significant difference between incubation with Pool 1 (fractions 1-5) or Pool 3 (fractions 9-10) from Cre or control mice, suggesting the absence of functional Cre exRNA. However, a significant increase in Nanoluc bioluminescence (\**p* < 0.05) was observed in FLEx reporter cells incubated with pool 2 (fractions 6-8) from Cre injected animals (84011 RLU) as compared to controls (72295 RLU), suggesting bdEVs can deliver functional Cre mRNA *ex vivo* (Figure 6G). To confirm our results, Cre activity was confirmed by RT-PCR of gDNA using discriminatory primer pairs between FLExON (recombined) and FLExOFF (non-recombined) gDNA. We detected a 50% increase (\**p* < 0.05) in pool 2-derived bdEVs from Cre injected animals compared to bdEVs of control injected animals (Figure 6H).

Collectively, our data provide evidence that the bdEVs enriched fraction from brain tissue are functionally active, being internalized by neurons and delivering functional Cre mRNA cargo with the ability to induce Cre activity in recipient cells.

## 4 DISCUSSION

In this study, we report a sensitive bioluminescence reporter system that allows to track the uptake of Cre species mediated by extracellular mechanisms, particularly bdEVs, by the permanent recombination at LoxP DNA sites. Upon establishing a striatal source of bdEVs carrying Cre mRNA in mouse brain, we showed a spatial gradient of tdTomato expression in the brain up to 3500µm away from the bdEV-donor cells and detected EVs containing Cre mRNA circulating in the bloodstream. Additionally, upon extracting bdEVs from striata of injected mice, we confirmed bdEVs morphology and integrity and observed the transfer of functional Cre mRNA from bdEVs to primary hippocampal neurons.

BdEVs detected in biofluids, such as serum and cerebrospinal fluid (CSF) have been studied as potential diagnostic and prognostic biomarkers for brain diseases (Badhwar and Haqqani 2020; Hill 2019; Rufino-Ramos et al. 2022; Street et al. 2012). However, the physiological role of bdEVs in brain communication to near and long distances remains largely unknown. Conclusions about their functions have been based on *in vitro* experiments, typically using a disproportionally high number of concentrated EVs exposed to a small population of neuronal cells for short period of time to increase the sensitivity of detection (Gupta, Zickler, and El Andaloussi 2021) and are thus not fully representative of physiological conditions *in vivo*. In this study, we aimed to translate previous *in vitro* findings to animal models by tracking the uptake of endogenous bdEVs carrying Cre mRNA in reporter cells through DNA recombination within the brain environment using floxed reporters in transgenic mice.

To create a localized brain region continuously secreting physiological levels of EVs carrying exogenous RNAs, we started by transducing the striatum of Flox-tdTomato Ai9 reporter mice through intracranial injection of a lentiviral vector encoding Fluc and Cre genes. These transgenes were regulated by ubiquitous promoters to ensure gene expression in the majority of cell types (Wettergren et al. 2012). To minimize the content of recombinant proteins in EVs, H2B and NLS peptides were fused to Fluc and Cre, respectively. Both proteins were predominantly located into the nucleus, while their mRNA products remained predominantly in the cytoplasm, thereby accessible to be loaded into EVs. Indeed, we detected Cre mRNA in EVs, even without having EV packaging signals, such as zipcodes (Bolukbasi et al. 2012) or exomotifs (Garcia-Martin et al. 2022; Shurtleff et al. 2016; Villarroya-Beltri et al. 2013), which suggests that Cre mRNA is naturally packaged into EVs, probably due to its overexpression in donor cells. On the contrary, we were not able to detect Cre protein in EVs, which is in line with other reports (Ridder et al. 2014; Steenbeek et al. 2018; Zomer et al. 2015, 2016). Even though the Cre protein remained undetected in EVs in our work, it has previously been described to be packaged in EVs through passive loading (Frühbeis et al. 2013) or by direct fusion with transmembrane proteins (Sterzenbach et al. 2017). Different findings can be potentially associated with the lack of nucleus targeting sequences fused to Cre, detection limit differences in RT-PCR and western blot, and EV isolation methods.

Our data demonstrates that long-term transduction of striata with lentiviral vectors encoding Cre and Fluc genes can be established in the brain and monitored by Fluc bioluminescence in living animals, suggesting a permanent neural source of Cre bdEVs was achieved in a limited brain region. Indeed, we isolated bdEVs from brain tissue of these animals and investigated the delivery of functional mRNA *in vitro*. Our optimized bdEV isolation protocol using a 4-layer ODG provided the isolation and enrichment of bdEVs restricted to just 3 fractions as confirmed by the presence of typical EV-markers and bdEV-cupped shaped morphology. Of note, despite bdEVs enrichment based on traditional EVs markers, including HSC70 and Flotilin1, further optimization would be useful to reduce cell-derived contaminants, similarly to what was previously described in (Huang et al. 2020). In this study, we carried out DNAse treatment prior to ODG to reduce nucleic acids contaminants that might lead to aggregation of bdEVs, proteins and cell contaminants. However, further measures can be taken to eliminate the co-isolation of DNA originating from nuclei disruption or organelle isolation, such as amphisomes or autophagosome (Fader et al. 2008; Jeppesen et al. 2019), adsorption of corona protein contaminants on bdEVs surface (Tóth et al. 2021) or the co-isolation of particles with overlapping size and density either biologically or protocol driven. This protocol could be improved by increasing the time of tissue digestion from 20 minutes to several hours together with a reduction of manual disruption and immunocapture of bdEVs based on surface markers.

Although ubiquitous promoters were used, our study was mainly focused on bdEVs secreted by neurons and partially astrocytes, as oligodendrocytes and microglia were not significantly transduced by the lentivirus vector. Interestingly, we also observed diffusion of bdEVs from the brain to other body compartments. Cre mRNA was found associated with serum circulating bdEVs after isolation by SEC and tetraspanin immunoprecipitation based on CD63, CD81 and CD9 presence. BdEVs isolated from transduced neural cells were not only enriched in Cre mRNA, but also retained their integrity, functionality and capacity to deliver Cre mRNA to recipient cells, corroborating their role in transferring functional cargo within the brain compartment and beyond. The analysis of tdTomato expression in peripheral organs would also be informative to evaluate to what extent bdEVs can deliver functional cargo beyond the CNS. Additionally, to confirm peripheral diffusion of bdEVs secreted from neural cells into the bloodstream, future analyses should focus on methods avoiding direct cell transduction to prevent transduction of other cell types present in the brain (Rufino-Ramos et al. 2022). Definitive surface markers for bdEVs would be helpful to distinguish different subpopulations of neural EVs, as neuronal markers such as L1CAM or NCAM were previously shown to be present in EVs from other tissues (Norman et al. 2021; Ter-Ovanesyan et al. 2021) and are thus not exclusive of neural cells. Indeed, it has recently been reported that neural cell type-specific EV markers exist for excitatory neurons (ATP1A3, NCAM1), astrocytes (LRP1, ITGA6), microglia-like cells (ITGAM,LCP1), and oligodendrocyte-like cells (LAMP2, FTH1) (You et al. 2022).

To detect the transfer of functional cargo by bdEVs at physiological levels, we used a reporter system encoding for an inverted sequence of the Nanoluc gene between Lox regions. Due to its high sensitivity, the expression of Nanoluc in reporter cells is beneficial to detect a low number of Cre-mediated functional events in a limited timeframe. Indeed, our previously published FLExNanoluc system (Breyne et al. 2022) showed robustness in detecting Cre activity mediated by EV delivery through the expression of Nanoluc. In the brain, we took advantage of the well-established Flox-TdTomato Ai9 mouse model (Madisen et al. 2010), that expresses tdTomato upon Cre recombination, to track the uptake of bdEVs carrying functional Cre mRNA. To allow the detection of tdTomato expression distally from the injection site, we narrowed the region of interest by performing brain coronal sectioning prior to analysis. Indeed, tdTomato expression was demonstrated following intracranial injection of concentrated EVs containing Cre mRNA in the striatum of Ai9 mice, overcoming the need for additional steps, such as fluorescent cell sorting of brain cells (Abels, Broekman, et al. 2019; Patel et al. 2016; Ridder et al. 2015; Zomer et al. 2015).

Lentiviral vectors do not replicate and are mostly localized in a restricted region surrounding the injection site as compared to an intracranial injection of AAV vectors (Parr-Brownlie et al. 2015). The continuous secretion of bdEVs carrying Cre mRNA from the injection site to the surrounding areas allowed tracking of bdEV-mediated communication to other Ai9 neural cells in physiological conditions overtime. The uptake of bdEVs containing functional Cre mRNA induced a permanent DNA recombination in recipient Ai9 neural cells was detected by ddPCR for tdTomato mRNA in brain sections. This technique enabled us to reveal the distribution patterns of bdEVs produced and secreted by neurons and astrocytes in the striatum. Of note, lentiviral vector expression was mainly restricted to the injection site since they are highly fusogenic and unable to replicate *in vivo*. We observed a spatial gradient of tdTomato expression from the injection site into the rostral and caudal regions, caused by the continuous spreading of functional Cre exRNA, with a peak at the injection site (0 µm) primarily caused by the lentiviral injection and magnified by bdEVs diffusion at short distances in the brain overtime. A previous study, detected a similar spatial gradient 500 µm away from the injection site after an intracranial injection of AAV8 encoding Cre into a CD63-floxed mice, leading to the secretion of CD63-GFP protein in bdEVs from the injection site to the surrounding regions (Men et al. 2019). Surprisingly, we were able to detect tdTomato expression 3500 µm away from the injection site, possibly due to permanent DNA recombination following the uptake of EVs carrying Cre mRNA in Ai9 reporter cells. Although both methods were able to detect long-term spreading of bdEVs in brain cells, further optimization should be considered to distinguish primary bdEVs transporting functional Cre molecules and secondary bdEVs transporting the product of Cre recombination within the brain. The methodologies used in both studies could not overcome or distinguish the potential spreading of reporter coding forms in bdEVs within the brain, including the spreading of tdTomato exRNA in bdEVs or CD63-GFP protein bdEVs from the injection site to other regions, respectively. Despite that, differences in signal intensity coming from primary or secondary bdEVs should exist since primary Cre-mediated recombination may result in higher expression levels rather than transfer of secondary product mRNA or proteins. We attempted to overcome this obstacle by showing communication between striatum secreting bdEVs carrying Cre mRNA and cerebellum containing the reporter system, even though we did not achieve reliable success (Supplementary Figures 5D, 5E and 5F). To overcome this issue, future analysis of the recombined sequence at genomic DNA level would dismiss the contribution of secondary bdEVs transporting tdTomato molecules. Moreover, other types of extracellular communication recently described such as exomeres, supermeres and tunneling nanotubes (Khattar et al. 2022) cannot be discarded and could account for the spreading of both Cre forms and secondary reporters.

In conclusion, our work demonstrates active brain communication between neural cells through bdEVs. Cre-LoxP systems allow the detection and permanent recording at DNA level of the uptake of physiological levels of bdEVs. BdEVs mediated the delivery of functional Cre mRNA to distal brain regions *in vivo* and *in vitro* thus results in genomic footprints in recipient cells. By mimicking the continuous physiological secretion of Cre exRNA in the brain we were able to corroborate previous *in vitro* findings and provide further evidence for functional bdEV delivery *in vivo* and *in vitro*. The spatio-temporal control of both source cells secreting Cre-containing bdEVs and LoxP reporter systems within the brain will contribute to revealing the role of bdEVs in extracellular communication.

## Acknowledgements

We thank Shilpa Prabhakar, Sevda Lule and Lisa Nieland for helping in bioluminescence acquisition. We thank João Peça and Joana Guedes for the support with Ai9 colony. We thank Ana Luisa Cardoso for the support with MACs system. CMV-STEAP3-SDC4-NadB booster plasmid and CD63-Nanoluc was kindly provided by Martin Fussenegger. We thank M. Zuzarte and LABCAR (Faculty of Medicine, University of Coimbra) for electron microscopy imaging. We thank Luisa Cortes, Margarida Caldeira, Tatiana Catarino and the CNC MICC team for assistance with microscopy imaging and Imaris processing. We thank Elisa Corti, Ivan Lalanda and Carlos Duarte for assistance with primary cell culture. We thank all members of the L.P.d A. and X.O.B. labs for all the support, discussion and comments. Schematic figures were created using Biorender.com.

## Author contribution

D.R.R., K.L., X.O.B., K.B. and L.P.A. conceived and designed the experiments. D.R.R., K.L., and M.P., T.V.S., and S.M. performed the experiments. D.R.R., K.L., K.O.B., P.R.L.P., M.P., M.S., T.V.S., S.M. and K.B. analyzed the data. D.R.R., K.L., K.O.B., P.R.L.P., M.P., M.S., T.V.S., S.M., X.O.B., K.B. and L.P.A. wrote and edited the paper.

## Methods

### Animals

C57BL/6 and BALB/c mice (Charles River Laboratories) were maintained in groups (2–5 per cage) in plastic cages (365 × 207 × 140 mm) with unlimited access to water and food under a 12-hour light/dark cycle at a room with constant temperature (22 ± 2 °C) and humidity (55 ± 15%). Equal number of male and female mice ranging from 8-10 weeks in age were randomly assigned to experimental groups. Animals were allowed 1 week of acclimatization to the surroundings before the beginning of stereotaxic injections. Physical state of animals was evaluated daily, and weight measured every week.

All animal experimental protocols were approved by: the Massachusetts General Hospital Institutional Animal Care and the European Union Directive 86/609/EEC for the care and use of laboratory animals. This study is part of a research project which was approved by the Center for Neuroscience and Cell Biology ethics committee (ORBEA_66_2015_/22062015 and ORBEA_289_) and the Portuguese Authority responsible for the regulation of animal experimentation, Direcção Geral da Agricultura e Veterinária (DGAV 0421/000/000/2015).

Researchers received adequate training (Federation of European Laboratory Animal Science Associations (FELASA)-certified course) and certification from Portuguese authorities (Direcção Geral de Alimentação e Veterinária) to perform the experiments.

### Lentiviral production, isolation and titer assessment

Lentiviral vectors encoding for the PGK-Cre-UbC-Fluc plasmid and FlexNanoluc plasmid were produced in the human embryonic kidney 293 (HEK293T) cell line with a three-plasmid system, following Addgene recommendations. Briefly, cells were seeded and 24h later, transfected with psPAX2 (Addgene plasmid #12260) and pMD2.G (Addgene plasmid #12259) packaging plasmids and CreFluc or FlexNanoluc plasmids. Six hours after transfection, cells were washed with PBS and incubated in new culture media. Lentiviral vector isolation was performed 48h-72h later upon ultracentrifugation at 70,000g followed by pellet re-suspension in 1% PBS/BSA. The viral particle content was evaluated by assessing HIV-1 p24 antigen levels by ELISA 2.0 (Retro Tek, 0801002), in accordance with the manufacturer’s instructions. Concentrated viral stocks were stored at −80 °C until use.

### Stereotaxic injection into the mouse brain

C57BL/6J mice (8-10 weeks of age) were anesthetized using 2.5% isoflurane in 100% oxygen via a nose cone. Mice were stereotaxically injected into the striatum with the following coordinates: anteroposterior 0.6 mm, lateral: ±1.8 mm, ventral: 3.3 mm relative to bregma and tooth bar: 0, with concentrated lentiviral vectors in a final volume of 3μl/injection containing 400 ng p24 antigen (capsid protein). Control animals were injected with 3μL 1% PBS/BSA. For cerebellar injections in deep cerebellar nuclei (DCNs) we used the following coordinates: anteroposterior: -6.5mm, lateral: ±0.75mm, ventral: -3.3mm relative to bregma (bregma and lambda aligned). Lentiviral vectors were injected in a final volume of 4μl/injection containing 450 ng p24 antigen. The infusion was performed at an injection rate of 0.25 mL/minutes using a 10 mL Hamilton syringe, 5 minutes after the infusion was completed, the needle was retracted 0.3 mm and allowed to remain in place for an additional 3 minutes prior to its complete removal from the mouse brains (Carmona et al. 2017). The skin was closed using a 6-0 Prolene® suture (Ethicon, Johnson and Johnson, Brussels, Belgium). Mice were kept in their home cages for the corresponding experimental period, before being sacrificed for EVs’ enrichment, western blot, qPCR and immunohistochemical analysis.

### *In vivo* bioluminescence analysis

Stable lentiviral transduction in the brain was monitored by assessing firefly luciferase bioluminescence periodically, using a Xenogen IVIS 200 Imaging System (PerkinElmer). For each determination, mice were anesthetized using 2.5% isoflurane in 100% oxygen via a nose cone and injected IP with D-luciferin (100 mg/kg). Bioluminescence images were acquired 5-10 minutes after D-luciferin injection. Analysis was performed using Living Image software 4.3.1 (PerkinElmer) and quantification of the bioluminescent signal was obtained from a region of interest (ROI) drawn around the cranium.

### Mouse tissue preparation for immunohistochemistry and DNA/RNA extraction

One to four months after lentiviral vector injections mice were perfused with 1% PBS under lethal administration of Ketamine and Xylazine injected IP. Blood and brain were collected and stored at -80°C. Mouse brains were coronally sectioned with 16 μm thickness on a freezing cryostat (Leica Microsystems, CM3050S). Brain sections were alternately collected for immunohistochemistry or RNA/DNA extraction.

### Immunohistochemistry

Brain sections were post-fixed with 4% PFA for 10 minutes and then washed with PBS three times and incubated 30 minutes with blocking solution (PBS/0.1% Triton X-100 containing 10% normal goat serum (Sigma-Aldrich)) and then incubation overnight at 4°C in blocking solution with primary antibodies: mouse anti-CRE (Sigma, F3165-2MG, 1:1000); mouse anti-Luciferase (Sigma, F3165-2MG, 1:1000); mouse anti-Parvalbumin (Sigma, F3165-2MG, 1:1000); mouse anti-MAP2 (Sigma, F3165-2MG, 1:1000); rabbit anti-NeuN (Sigma, F3165-2MG, 1:1000); rabbit anti-IBA1 (Sigma, F3165-2MG, 1:1000). Sections were washed with PBS and incubated for 2h at RT with the secondary antibodies: goat anti-mouse IgG Alexa Fluor 488 (Thermo Fisher, A31560, 1:500) and goat anti-rabbit IgG Alexa Fluor 647 (Invitrogen, A32728, 1:1000) diluted in blocking solution. The sections were washed with PBS and incubated during 10 minutes with DAPI (1:5,000; Sigma), washed, and mounted with Vectashield Antifade Mounting Medium (Vector Labs, H-1000). Immunofluorescence was visualized and imaged with a Keyence BZ-X810 microscope, a Zeiss LSM 510 Meta confocal microscope (Carl Zeiss MicroImaging), equipped with EC Plan-Neofluar 40x/1.30 Oil DIC M27 (420462-9900) and Plan-Apochromat 63x/1.40 Oil DIC M27 (420782-9900) objectives and LSM Image software.

### Imaris 3D rendering

Carl Zeiss z-stack laser scanning confocal image files were reconstructed using Imaris software (Bitplane, version 9.6.1). The phalloidin staining was used to create a 3D cell-surface mask (represented in pink) that was then applied to select the brain-derived EVs (green dots) present inside of the neuronal cells.

### Cell culture and transduction

HEK293T cells were maintained in standard Dulbecco’s Modified Eagle’ medium (DMEM; Sigma) supplemented with 10% fetal bovine serum (Life Technologies) and 1% penicillin/streptomycin (Gibco) and grown at 37 °C in 5% CO2. Stock cells were passaged 2–3 times/week with 1:6 split ratio and used within 10-20 passages. Cells were tested for mycoplasma contamination monthly and found negative. Cells grown for EV isolation were cultured in media supplemented with 10% EV-depleted FBS (FBS was depleted of EVs by 18 h centrifugation at 100,000 × g and the resulting supernatant was filtered at 220nm).

### Cell transduction

HEK293T cells were transduced 24h after plating with lentivirus vectors encoding PGK-Cre-UbC-Fluc or FlexNanoluc constructs at a ratio of 400 ng p24 antigen per 200,000 cells. Twenty-four hours later, the medium was replaced with regular medium and cells were cultured and expanded under standard conditions. Luminescence (PGK-Cre-UbC-Fluc construct) and fluorescence (FlexNanoluc construct) were monitored weekly.

### Transwell of Cre and FlexNanoluc expressing HEK 293T cells

HEK 293T cells transduced with the FlexNanoluc construct were seeded in the bottom chamber of 12-well plates at 100,000 cells/well in DMEM (Thermo Fisher) supplemented with 10% FBS (Thermo Fisher). Meanwhile, HEK 293T cells transduced with the Cre construct were seeded in the upper chamber of a 1.0-μm-pore Transwell system at 50,000 cells/well in DMEM supplemented with 10% FBS. After 24h, Cre expressing cells were transfected with the CMV-STEAP3-SDC4-NadB plasmid (Dautzenberg, Rabelink, and Hoeben 2021) to boost EV production. Control Cre cells were not transfected. Six hours following transfection, cells were washed in PBS and fresh media was added. Twelve hours later, the transwell systems seeded with Cre expressing cells were incubated with FlexNanoluc expressing cells in 12-well plates. After 48h, cells from the bottom chamber were collected with Passive Lysis Buffer (Promega), luminescence was measured and DNA extraction was performed as described elsewhere.

### Bioluminescence assays

Firefly luciferase and Nanoluc expression in EVs, cells and cerebellum collected with Passive Lysis Buffer (Promega), were analyzed with the addition of luciferin (100mg/mL) or furimazine (Nano-Glo® Luciferase, Promega) diluted 1:200 to 1:500 in 1X PBS, respectively. Samples were incubated with the reagent for at least 1 minutes prior to reading on Synergy H1 Hybrid Multi-Mode Reader (BioTek) or FLUOstar Omega Microplate Reader (BMG LABTECH). At least two reads were performed on each sample, and the average values were considered for analysis. For luminescence readings, samples were loaded into white 96-well culture plates (Lumitrac 200) or opaque 96-well plate (Corning). Each sample was loaded in duplicate with a volume of ranging from 20 to 100 µL in each well.

### Isolation of EVs by Size Exclusion Chromatography (SEC)

Conditioned medium was collected from cells after 48-72h (approximately 80% confluency) and centrifuged at 300 x g for 5 minutes to remove cellular debris. The supernatant was then concentrated with 100 kilodalton (kDa) molecular weight concentrator (UFC9100, Amicon® Ultra-15 Centrifugal filters) to a final volume of 0.5 ml (spin at 6,000 x g for 15 minutes). Concentrated media was loaded onto qEV Original SEC columns (SP1, IZON Science) and 500 µL fractions were collected by elution with PBS using the Automatic Fraction Collector (AFC) according to the manufacturer’s protocol. The first 5 fractions correspond to High particle/low protein fractions (typically described as fractions from 7 to 11) were further concentrated with 30 kilodalton (kDa) molecular weight concentrators (UFC503096, Amicon®Ultra-0.5 Centrifugal filters) to a final volume of 50 to 100 µL.

### EVs enrichment from brain tissue

A thick coronal section (1-2 cm of thickness) from the injection site was collected per mouse and stored at -80°C, until further processing. The frozen tissue was sliced lengthwise on ice to generate 1–2 cm long, 2–3 mm wide tissue sections (Huang et al. 2020; Su et al. 2021; Vella et al. 2017). The tissue pieces from each sample were weighed and incubated with 50 U/mL collagenase type 3 (#CLS-3, CAT#LS004182, Worthington) in Hibernate-E medium (at ratio of 8μL/mg tissue) in a shaking incubator (25-27°C for 20 minutes). After 10 minutes of incubation samples were inverted twice, 5 minutes later pipetted up and down twice and incubated for another 5 minutes, followed by addition of ice-cold 10x inhibition buffer containing 10x protease inhibitors (cOmpleteTM Mini proteinase inhibitor (Roche), phenylmethylsulfonyl fluoride) and 10x phosphatase inhibitors (sodium orthovanadate and sodium fluoride) in PBS with a final concentration of 1x. The digested brain extracts were subjected to centrifugation step at 4°C, 300×g for 5 min. The supernatant was collected and centrifuged at 4°C, 2000×g for 10 minutes. The resulting supernatant was collected and further centrifugated at 4°C, 10,000×g for 30 minutes. 1mL Supernatant was then incubated with 5 µl of DNase (Sigma D-5025) 10mg/mL for 10 minutes and then filtered with 0.22 µm filter (Millipore).

The 10,000 × g supernatant was loaded on top of a 4-layer iodixanol density gradient (ODG) containing 40 mM, 20 mM, 10 mM and 5 mM OptiPrep reagent (Sigma-Aldrich) in ultra-clear SW41Ti tubes (Beckman Coulter). The iodixanol density gradients were centrifuged at 100,000 x g at 4◦C for 18 hours in SW41Ti rotor (Beckman Coulter). Ten fractions (F1, F2, F3, F4, F5, F6, F7, F8, F9, F10 each of 1 mL) were collected, weighed and densities calculated. Each fraction was subjected to a washing step in ice-cold PBS at 100,000 × g at 4°C for 2 h using a SW28Ti rotor (Beckman Coulter). The pelleted EVs were resuspended in ice-cold PBS. Samples were analyzed by NTA and then processed with AllPrep DNA/RNA/Protein Mini Kit (cat. no. 80004, Qiagen).

### Immunomagnetic isolation of EVs from serum

Up to 2mL of EVs isolated by SEC from serum of C57BL/6J mice stereotaxically injected in the striatum with lentiviral vectors encoding for Cre construct were incubated with 25uL of CD9, CD63 or CD81 MicroBeads (Miltenyi Biotec) overnight at 4°C in a tube rotator in the absence of light. Equilibration buffer (100µL) was applied on top of a µColumn (Miltenyi Biotec) that was previously placed in the magnetic field of the µMACS Separator attached to the MACS MultiStand and rinsed 3 times with 100µL of Isolation Buffer. The magnetically labelled samples were applied to the column which was placed in a mMACS Separator (Miltenyi Biotec). The column was washed 4x with Isolation Buffer and then placed in 1.5mL tubes. The sample was eluted by adding 100µL RNA lysis buffer (Miltenyi Biotec) to the column and flushed out by firmly pushing the plunger into the column. Downstream isolation of EV-derived RNA was performed using Total RNA Purification Plus Kit (Norgen) and according to manufacturer’s instructions. cDNA synthesis for mRNA was performed with iScript cDNA Synthesis Kit (Bio-Rad) and RT-PCR was performed with the Sso Advanced SYBR Green Supermix Kit (Bio-Rad) using the StepOnePlus Real-Time PCR System (Applied Biosystems).

### DNA, RNA and protein extraction

DNA, RNA and protein extractions were performed from cultured cells, brain sections and EVs following the protocol recommendations of the RNeasy Plus Micro Kit (cat. no. 74034, Qiagen) and AllPrep DNA/RNA/Protein Mini Kit (cat. no. 80004, Qiagen). Isolated DNA and RNA samples were quantified by Nanodrop (ThermoFischer Scientific) and Bioanalyzer 2100 (Agilent Technologies, Santa Clara, CA). Protein concentration was determined by Bradford assay (Bio-Rad Laboratories) for protein extracted from cultured cells or brain sections and micro bicinchoninic acid (microBCA) for protein extracted from EVs according to the manufacturer’s instructions (Bio-Rad Laboratories).

### cDNA synthesis and RT-PCR

RNA samples were reverse transcribed using the SuperScript VILO cDNA Synthesis Kit (ThermoFisher Scientific) and iScript Selected cDNA Synthesis kit (Bio-Rad) according to manufacturer’s instructions and stored at -20ᵒC. RT-qPCR was performed using the primers described in Table 1. Gene expression was determined using the SYBR green protocol qPCR mix, as prepared following the manufacturing protocol of Power SYBR Green PCR Master Mix (Applied Biosystems, Beverly, MA) and with the SsoAdvanced SYBR Green Supermix Kit (Bio-Rad). qPCR was started with enzyme activation by heating at 95°C during 10 min, followed by 40 cycles of two steps: 95°C for 20 s, and 60°C for 1 min. To verify PCR specificity a melting curve was performed, with the following program: 95°C for 20 s, 60°C for 1 min, and 60°C–95°C with an increment of 0.3°C per 15 s. RT-PCR was performed using QuantStudio 3 PCR system (Applied Biosystems) or StepOnePlus Real-Time PCR System (Applied Biosystems).

**Table 1.**
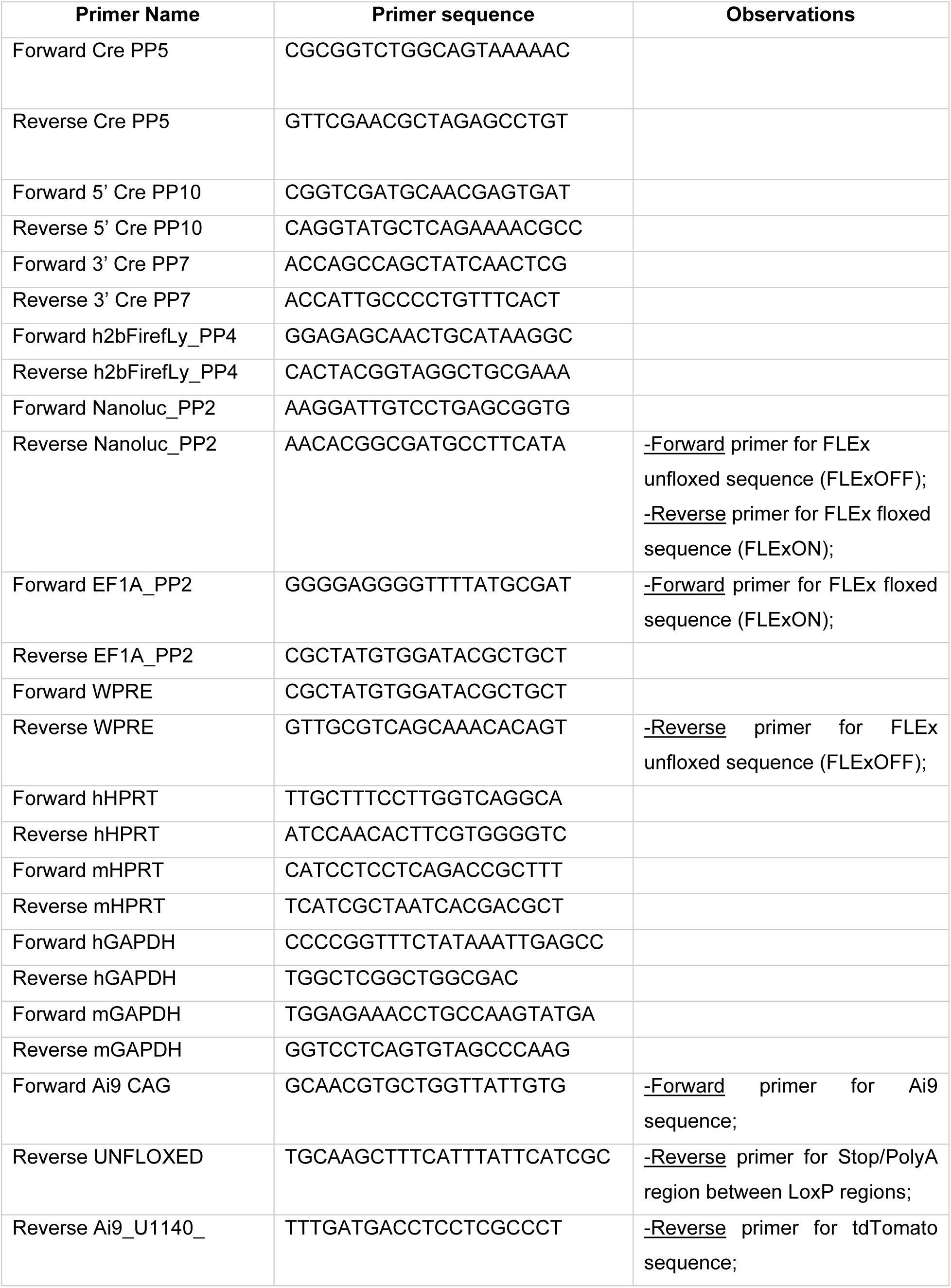
Primer sequences used in RT-qPCR.

### Digital droplet PCR (ddPCR)

To evaluate levels of gene expression of tdTomato and GAPDH in cells and brain coronal sections, gene expression of tdTomato [TaqMan probe FAM - Mr07319439_mr (Thermofisher)] and GAPDH (Rufino-ramos et al. 2022)(Al Ali et al. 2021) was analyzed using ddPCR following the PrimePCR ddPCR Gene Expression Probe Assay (Bio-Rad). Using the manufacturer’s protocol droplets were generated with DG8 Cartridge using a QX200 droplet generator and PCR was performed with thermal cycling conditions using QX200 Droplet Reader and QuantaSoft Software (Bio-Rad) to analyze mRNA levels.

### Western blotting

Total protein from cells and EVs was extracted in RIPA buffer [50 mM Tris-base; 150 mM NaCl; 5 mM EGTA; 1% Triton X-100; 0.5% sodium deoxycholate; 0.1% SDS supplemented with cOmplete Mini proteinase inhibitor (Roche) and 0.2 mM PMSF (phenylmethylsulphonyl fluoride), 1 mM DTT (dithiothreitol), 1 mM sodium orthovanadate and 5 mM sodium fluoride]. Protein concentration was determined by Bradford assay according to manufacturer’s instructions (Bio-Rad Laboratories). Protein samples were denatured (95 °C for 10 minutes) with 6×sample buffer [0.375 M Tris pH 6.8 (Sigma-Aldrich), 12% SDS (Sigma-Aldrich), 60% glycerol (Sigma-Aldrich), 0.6 M DTT (Sigma-Aldrich) and 0.06% bromophenol blue (Sigma-Aldrich)]. Samples were resolved by electrophoresis on 10 or 12% SDS-PAGE gels and transferred onto polyvinylidene fluoride (PVDF) membranes (GE Healthcare). Total protein labeling was performed using No Stain Labeling Reagent (Invitrogen) according to manufacturer’s protocol. Membranes were blocked by incubation in 5% non-fat milk powder in 0.1% Tween 20 in Tris buffered saline (TBS-T) and incubated overnight at 4°C with primary antibodies: ALIX (BD Biosciences, 611620, 1:1000), calnexin (Santa Cruz, sc-11397, 1:1000), CD63 (DSHB, AB528158, 1:500), CRE (Millipore, MAB3120, 1:1000), flotillin-1 (BD Biosciences, 610820, 1:1000), HSC70 (GeneTex, GTX101144, 1:1000), Lamp-2 (Santa Cruz, sc18822, 1:1000), TSG101 (BD Biosciences BD612696, 1:1000). Then, the membranes were washed 3 times in TBS-T for 10 minutes each and incubated with an alkaline phosphatase-linked secondary goat anti-mouse/anti-rabbit antibody (1:10,000; Thermo Scientific Pierce) at RT for 1h. Bands were visualized with Enhanced Chemifluorescence substrate (ECF) (GE Healthcare) in the chemifluorescence imaging (ChemiDoc Imaging System, Bio-Rad). Analysis was carried out based on the optical density of scanned membranes in ImageLab version 5.2.1; Bio-Rad.

### Characterization of EVs by Transmission electron microscopy (TEM)

Brain-derived EVs (isolated by ODG) were fixed with 2% PFA and allowed to absorb on Formvar-carbon coated grids (TAAB Laboratories) for 5 minutes. The excess liquid was blotted off the film surface using filter paper (Whatman). Then, the grids were contrasted with 2% uranyl acetate and after 1 minutes, the excess stain was blotted off and the sample air dried. Observations were carried out using a Tecnai G2 Spirit BioTwin electron microscope (FEI) at 100 kV.

### Characterization of EVs by Nanoparticle Tracking Analysis

Number of EVs diluted in PBS was assayed using Nanoparticle Tracking Analysis Version 2.2 Build 0375 instrument (NanoSight). Particles were measured by the acquisition of 5 videos of 30 s and the number of particles (30–800 nm) was determined using NTA Software 2.2. Samples were diluted 1:1000 in PBS prior to analysis. The following photographic conditions were used: frames processed (1498 of 1498 or 1499 of 1499); frames per second (24.97 or 24.98 f/s); calibration (190 nm/pixel); and detection threshold (6 or 7 multi). Number of particles per frame was within the recommended range of 20–100 particles/frame for NanoSight NS300.

### gDNA recombination analysis

To evaluate gDNA recombination mediated by Cre, 2 pairs of primers were generated to amplify either the floxed sequence or the unfloxed sequence (described in Table 1). Each pair of primers is either specific for the floxed sequence (gDNA recombination upon CRE activation) or the unfloxed sequence (non-recombined gDNA). Gene expression was determined using the SYBR green protocol qPCR mix, as prepared following the manufacturing protocol of Power SYBR Green PCR Master Mix (Applied Biosystems, Beverly, MA) and with the SsoAdvanced SYBR Green Supermix Kit (Bio-Rad). qPCR was performed using QuantStudio 3 PCR system (Applied Biosystems) or StepOnePlus Real-Time PCR System (Applied Biosystems).

### Primers (RT-PCR)

**Supplementary Figure 1.**
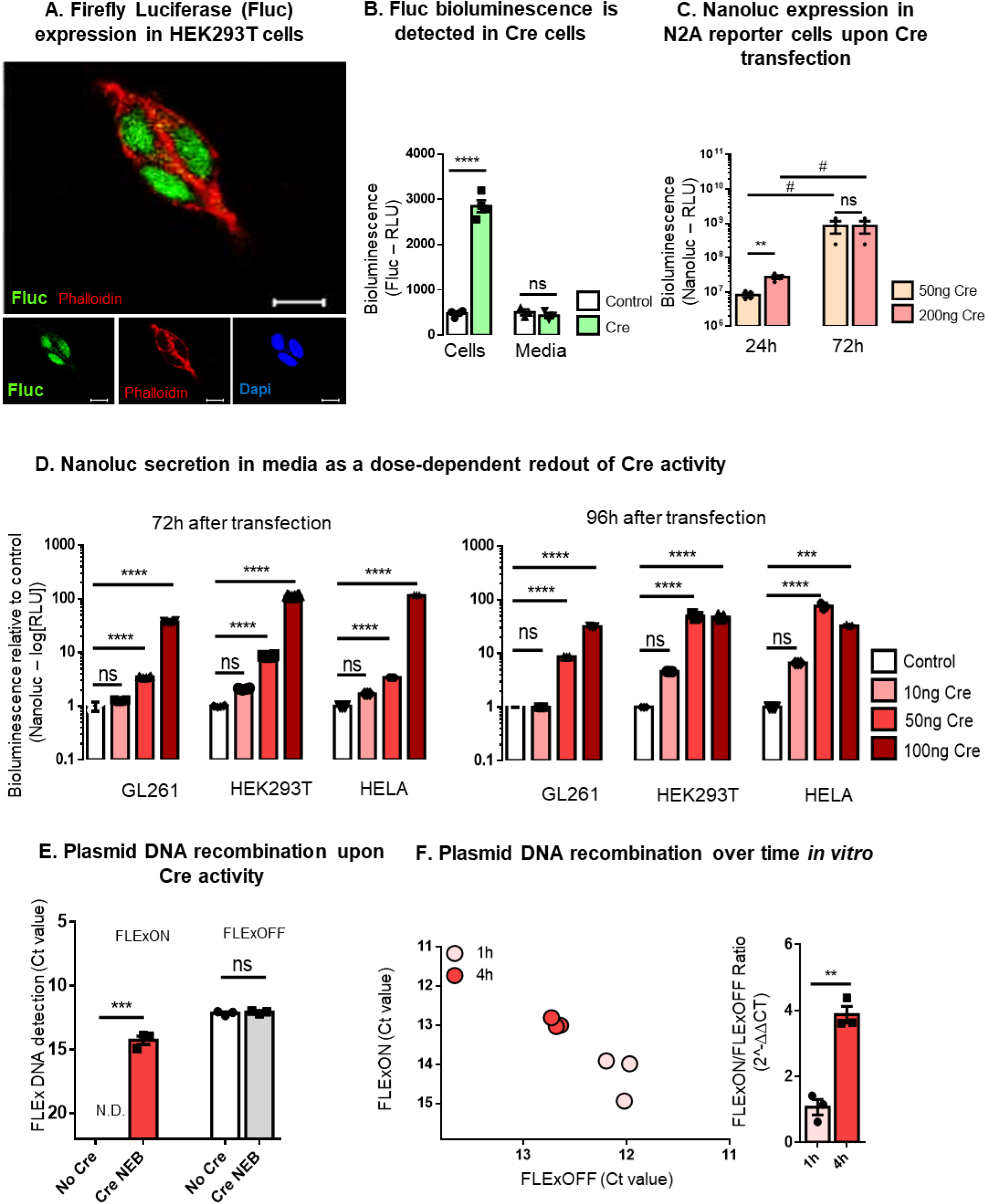
Firefly luciferase (Fluc) and Nanoluciferase (Nanoluc) bioluminescence as a tool to study extracellular communication. A. Representative immunofluorescent image from confocal microscopy of HEK293T cells stably expressing Fluc protein (green), mainly found in the nucleus (blue). Actin filaments in cytoplasm were stained with Phalloidin (red). Scale bar, 10 µm. B. Fluc bioluminescence was detected in cells expressing Cre, but not secreted into the media. Data presented as means ± SEM and compared by ordinary one-way ANOVA followed by Sidak’s multiple comparisons test (F=201.3). C. Bioluminescence evaluation upon co-transfection of Cre plasmid and FLExNanoluc plasmid into Neuro-2A cells was time dependent. Data presented as means ± SEM and compared by Unpaired t test (at 24h comparing 50ng to 200ng Cre plasmid/transfection) and Paired t test (comparing the same condition at 24h to 72h). D. Nanoluc secretion into media as a dose-dependent readout of Cre activity. GL261, HEK293T and HELA FLExNanoluc stable cell lines were generated and Nanoluc expression evaluated 72 and 96 h after transfection of Cre plasmid in three different doses (10ng, 50ng and 100ng). Data presented as means ± SEM and compared by ordinary one-way ANOVA followed by Dunnett’s multiple comparisons test. E. To evaluate plasmid recombination at the gDNA level, Cre recombinant protein (NEB biosciences) was incubated in a tube together with FLExNanoluc plasmid, recombination levels were evaluated using primer pairs designed to differentially detect FLExON and FLExOFF conditions. Data are presented as means ± SEM and compared by Unpaired t test. F. Under the same conditions as in E., plasmid DNA recombination was shown to increase from 1h incubation to 4 h incubation. Data is presented a mean ± SEM and compared by Unpaired t test. Statistical significance: #p < 0.05, *p < 0.05, **p < 0.01, ***p < 0.001, ****p < 0.0001 and ns for non-significant.

**Supplementary Figure 2.**
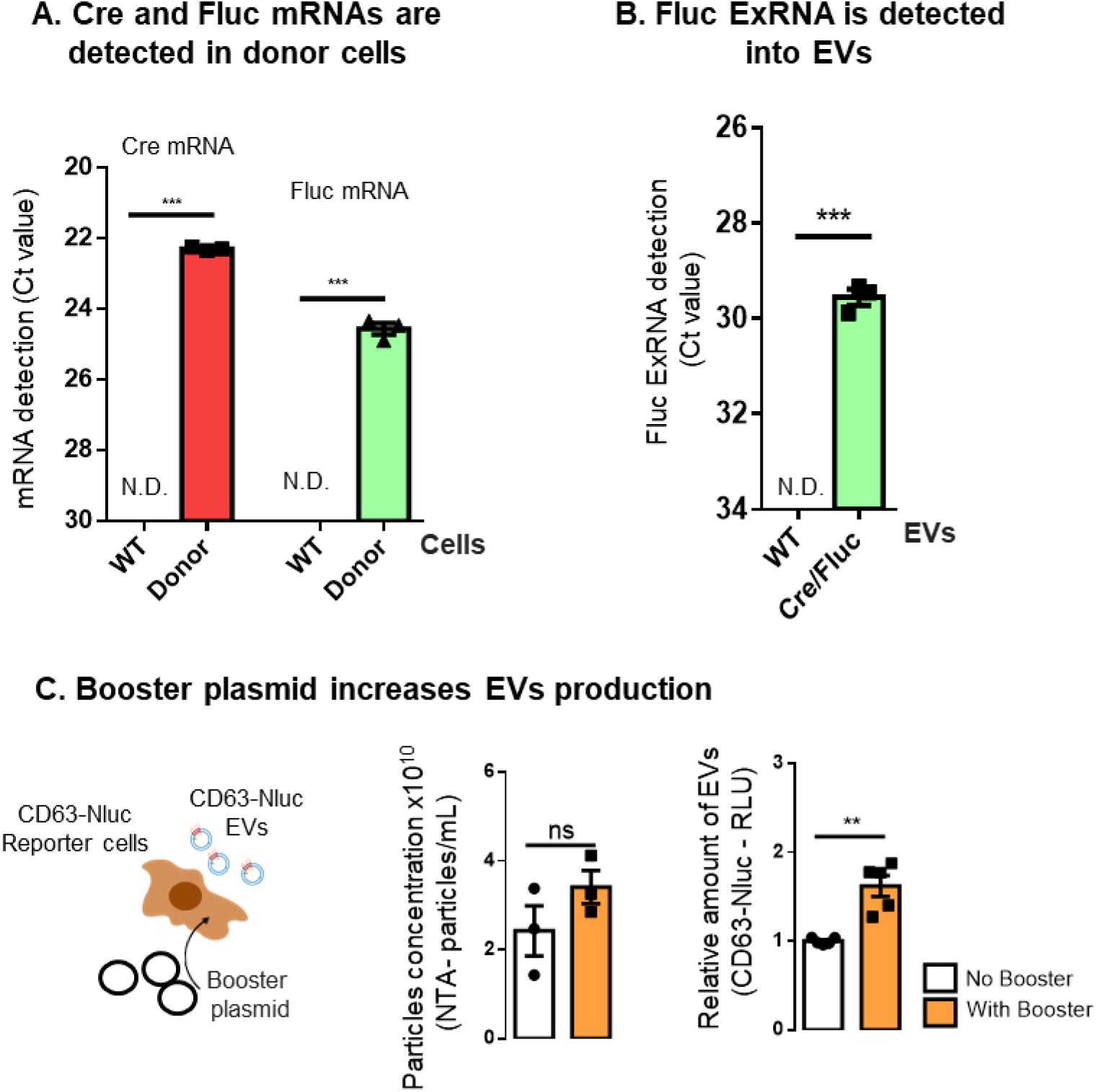
Detection of Cre and Fluc mRNAs in the system and boosting EV production. A. Cre and Fluc mRNAs were detected in stable transduced donor cells, but not in WT cells. B. Fluc exRNA was detected in EVs derived from Cre/Fluc cells, but not in EVs from WT cells. C. Tricistronic booster plasmid expressing CMV-STEAP3-SDC4-NadB (Kojima et al. 2018) was transfected into CD63-NanoLUC reporter cells to increase EV production. Difference between boosted and non-boosted CD63-Nanoluc EVs was evaluated for particles number by NTA and Nanoluc bioluminescence. Data is presented as means ± SEM and compared by Unpaired t test. Statistical significance: **p < 0.01, ***p < 0.001, and ns for non-significant.

**Supplementary Figure 3.**
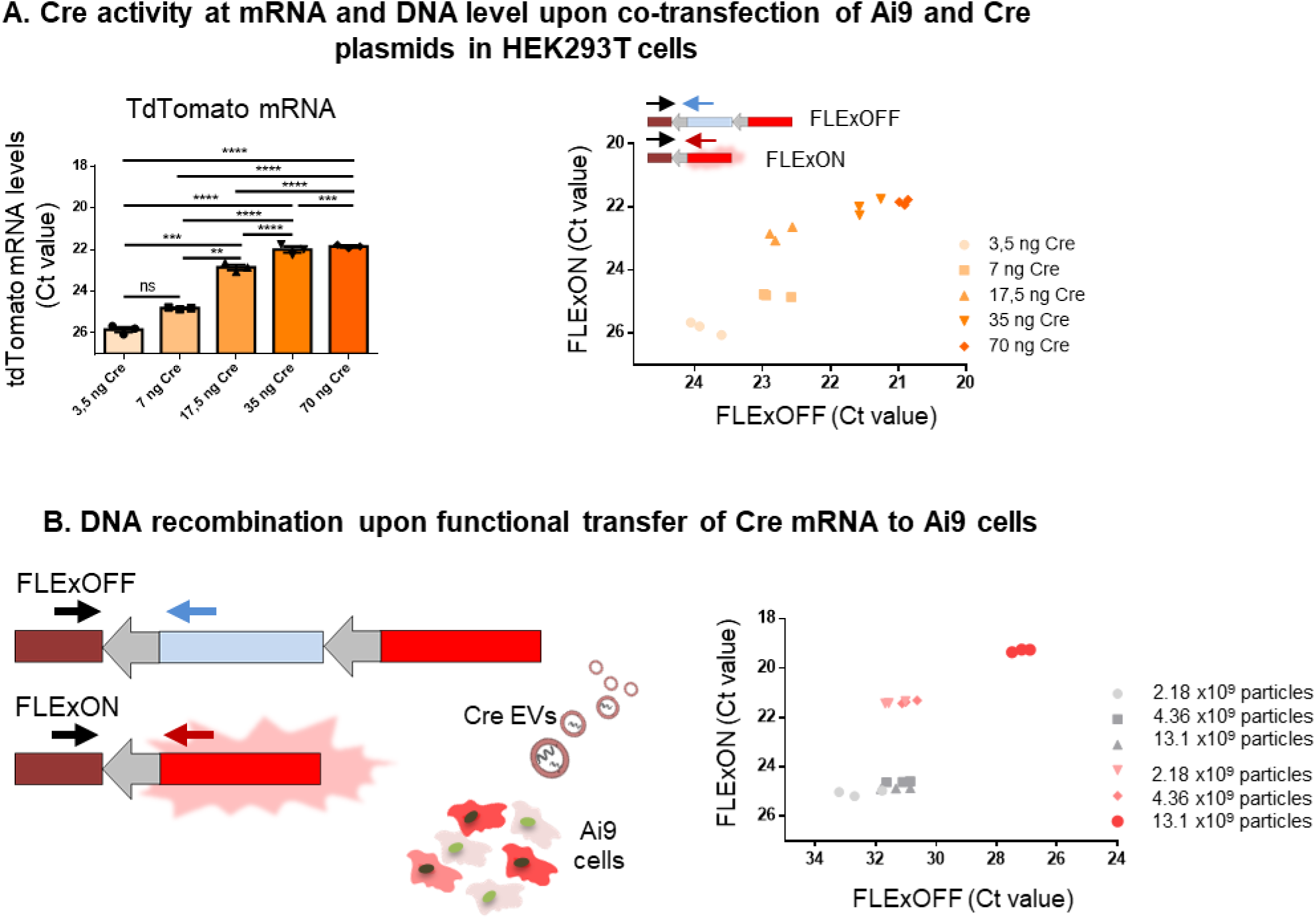
Evaluation of Cre activity in Ai9 reporter cells. A. Co-transfection of Ai9 and CRE plasmids (3.5ng, 7ng, 17.5ng, 35ng and 70ng) showed dose dependent expression of tdTomato mRNA in cells. The mRNA expression was then evaluated to distinguish FLExON (expression) and FLExOFF (no expression). Data presented as means ± SEM and compared by ordinary one-way ANOVA followed by Tukey’s multiple comparisons test (F = 293.0). B. The same system was used to distinguish FLExON (expression) and FLExOFF (no expression) at the DNA level after the cells had been incubated with 3 different doses of EVs carrying Cre mRNA (2.18, 4.36 and 13.1 X10^9^ particles. Statistical significance: *p < 0.05, **p < 0.01, ***p < 0.001, ****p < 0.0001 and ns for non-significant.

**Supplementary Figure 4.**
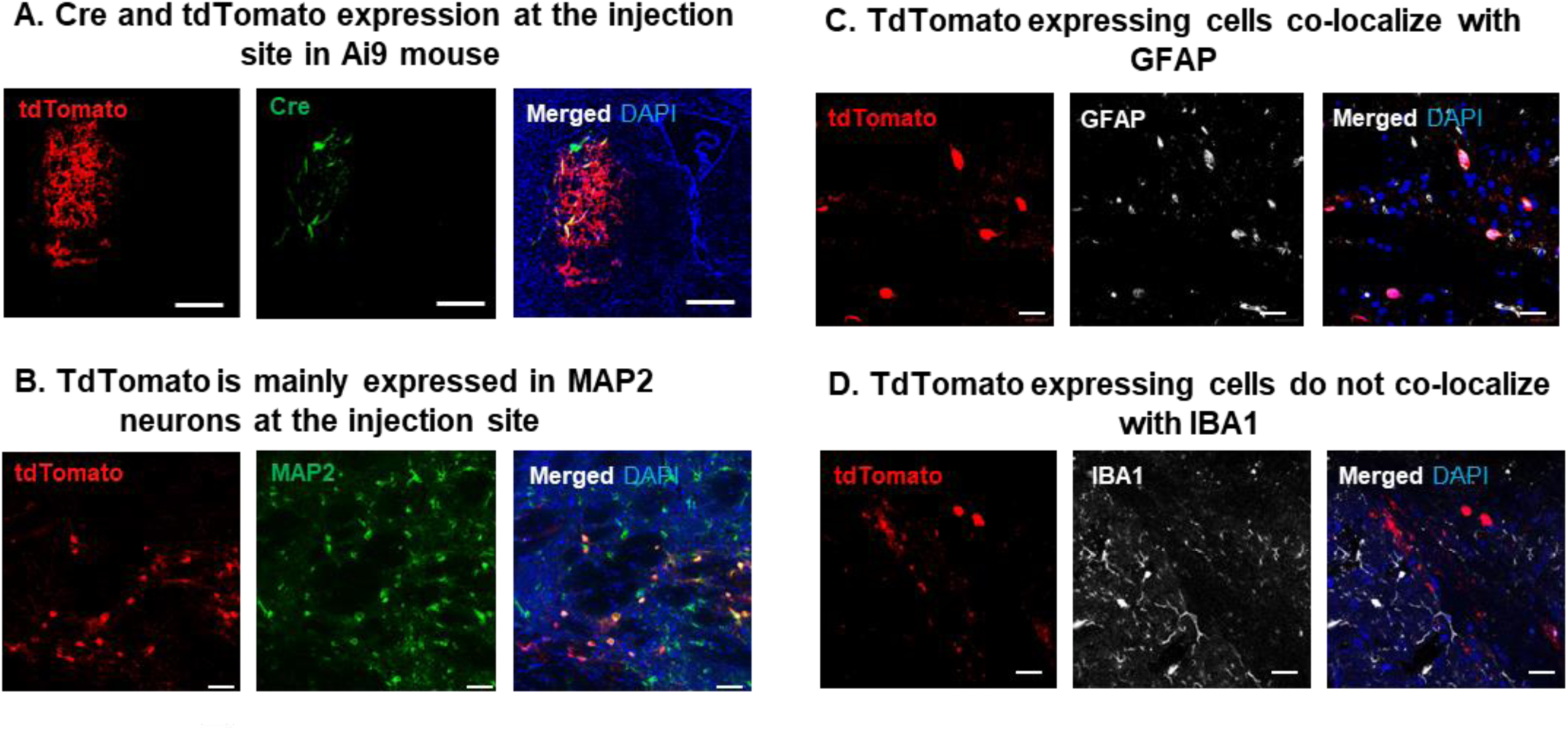
Transduced cells at the injection site in Ai9 mouse. A. Cre and tdTomato were mainly expressed at the injection site in Ai9 mouse. B. TdTomato positive cells co-localize with MAP2 neurons at the injection site. C. TdTomato positive cells partially co-localize with GFAP positive astrocytes. D. TdTomato positive cells do not co-localize with IBA1 positive microglia. Images are representative of a group of 5 Ai9 animals. Nucleus is represented by DAPI staining. Analysis performed with a Keyence BZ-X810 microscope 20x (Image A - injection site, scale bar 50μm) and laser confocal microscopy equipped with Plan-Apochromat 40×/1.40 Oil DIC M27 (420782-9900) (image B, C and D, scale bar 20μm).

**Supplementary Figure 5.**
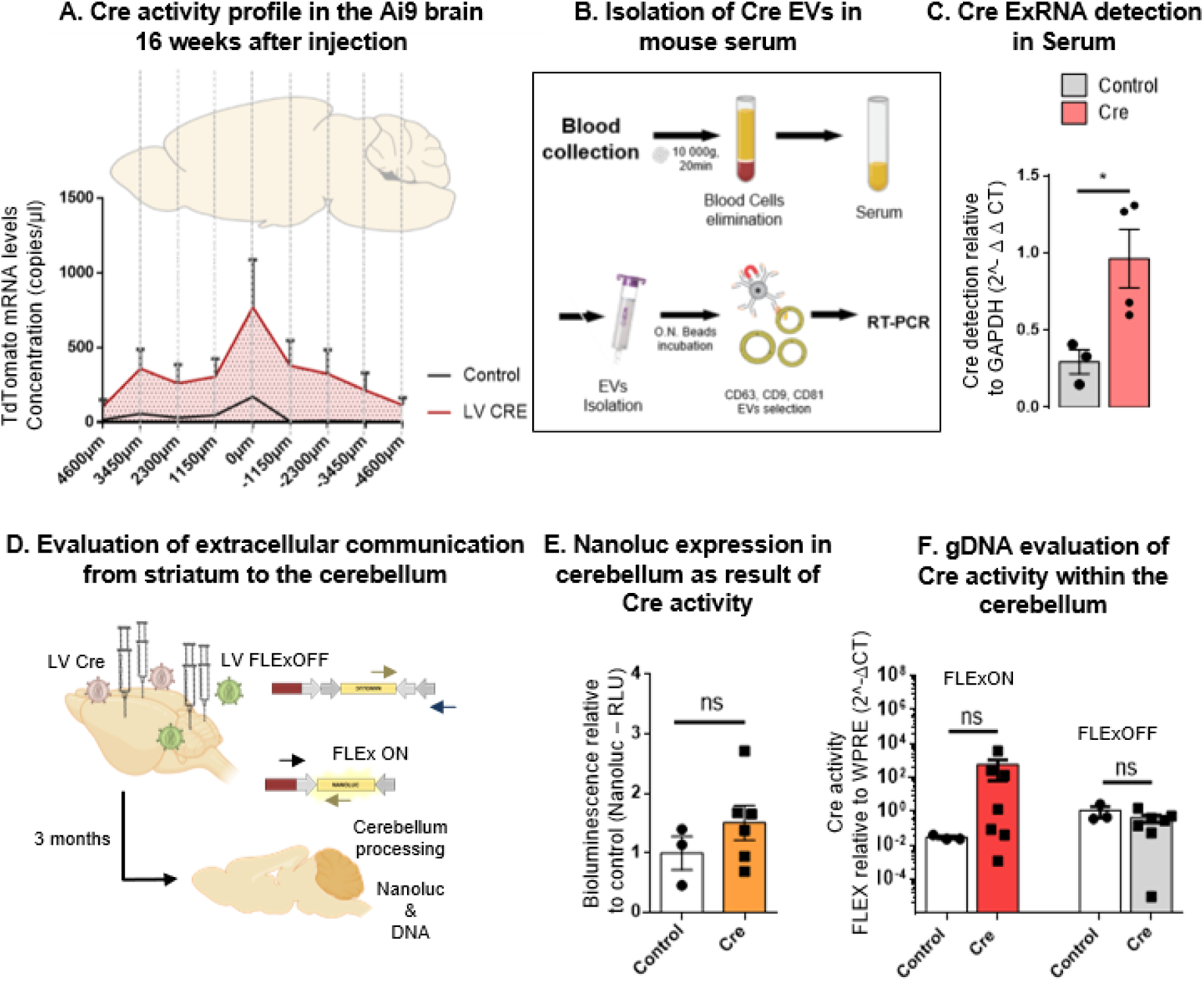
Extracellular communication demonstrated by Cre exRNA detection in the brain and periphery. A. Cre activity profile in the Ai9 mouse brain 16 weeks after injection. Whole-brain coronal sections were used to compare tdTomato mRNA expression levels in the brain of Ai9 mice injected with LV Cre (red) or 1%PBS/BSA (grey). B. Schematic illustration of the protocol used to isolate brain-derived EVs (bdEVs) from serum. Briefly, 1-2 mL of blood were collected at the time of sacrifice, then centrifuged at 10 000g for 20 minutes to remove blood cells and other cell particles. Serum was then concentrated using 100 kDa filters to a final volume of 500ul and then loaded onto qEV Original SEC columns. 5 fractions of 500uL (fractions 7 to 11) were collected and then incubated with MicroBeads recognizing the tetraspanin proteins - CD9 or CD63 or CD81 (MACS® Technology) overnight. RNA extraction was then performed on EVs bound to beads C. Detection of Cre exRNA in EVs collected from the serum of mice injected with 1%PBS/BSA (controls) and LV-Cre construct was evaluated by RT-PCR and normalized to GAPDH. Cre ExRNA was detected in tetraspanin positive EVs derived from LV-Cre injected mice when compared to control EVs (derived from non-injected mice) (N=3). Data were presented as means ± SEM and compared by one-way ANOVA followed by Dunnett’s multiple comparison test (F = 6.459), *p < 0.05. D. Schematic representation of double Cre sources created in the striatum by intracranial injection of LV Cre, and double reporter system created into the cerebellum upon injection of LV FLExNanoluc. Three months after injection animals were sacrificed and cerebella analyzed. E. Nanoluc Bioluminescence in the cerebellum was compared between controls and Cre treated group. F. gDNA levels in the cerebellum were evaluated in terms of FLExON (expression) and FLExOFF (no expression) between control and Cre injected animals. Data presented as means ± SEM and compared by Unpaired t test. Statistical significance: ns for non-significant.

**Supplementary Figure 6.**
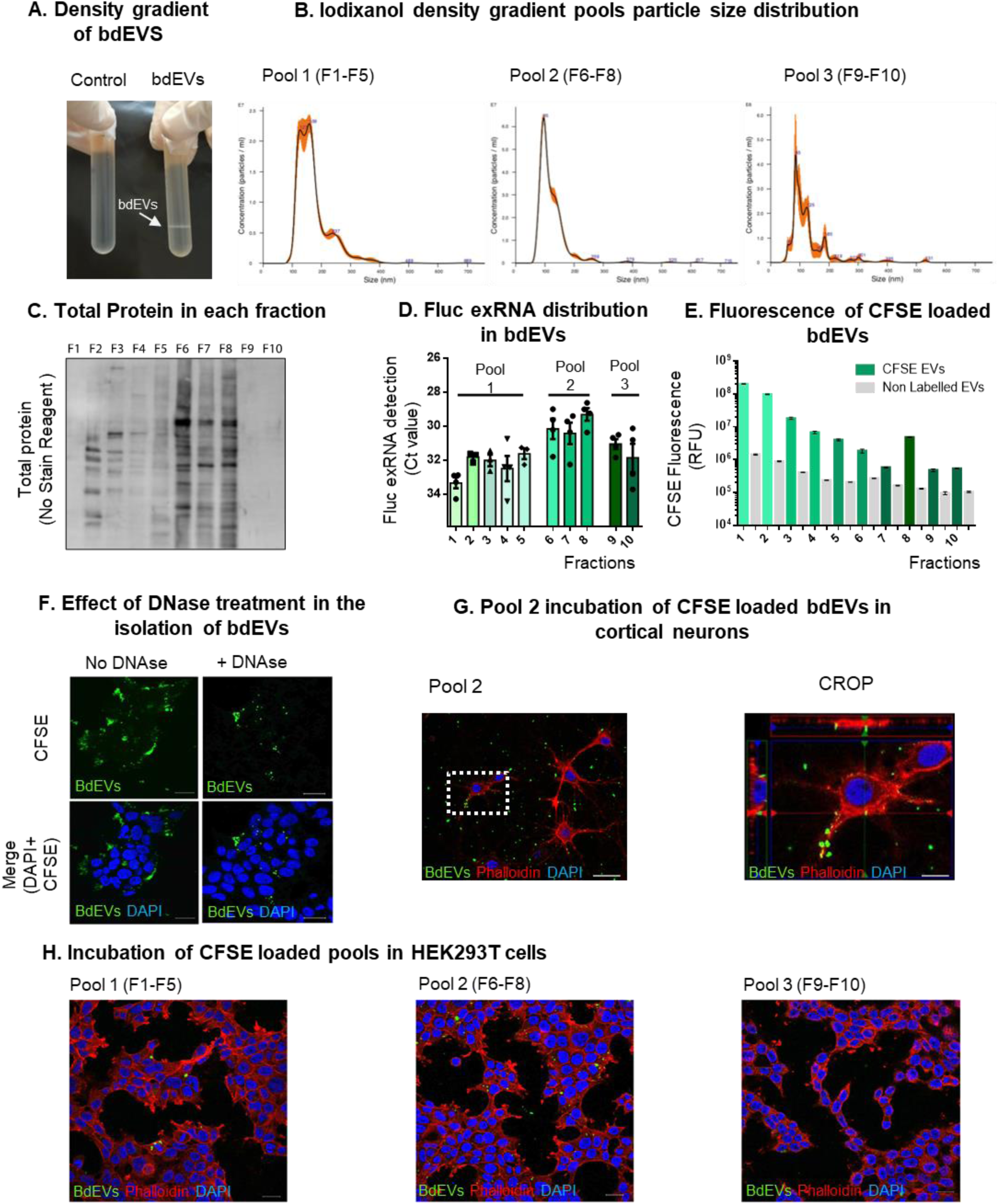
Characterization of brain derived EVs (BdEVs) A. Isolation of carboxyfluorescein succinimidyl ester (CFSE) bdEVs by density gradient separation. B. Size characterization by Nanoparticle tracking analysis (NTA) of pools of EVs, namely Pool1 (F1-F5), Pool2 (F6-F8) and Pool 3 (F9-F10). C. Total protein in each fraction. No Stain Labeling showed a high protein content in F6,F7 and F8. D. Fluc ExRNA detection in the 10 fractions collected after density gradient centrifugation. E. CFSE fluorescence distribution in the 10 fractions collected after density gradient centrifugation (green bars). Non-labelled control with all the fractions was used to detect the background fluorescence (grey bars). F. Effect of DNAse treatment in the sample before isolation of bdEVs by density gradient centrifugation. G. Pool 2 incubation of CFSE loaded bdEVs in primary hippocampal rat neurons for 6h. A high magnification image (crop) shows accumulation of brain-derived EVs internalized in specific cell compartments (green dots). F. Incubation of Pool 1, 2 and 3 of CFSE loaded EVs (green) in HEK293T cells for 6h. For experiment G and H cells were stained with phalloidin (red) and DAPI (blue) and analyzed by laser confocal microscopy equipped with Plan-Apochromat 40×/1.40 Oil DIC M27 (420782-9900). Scale bar 20 μm and 5 μm (crop).

## References

Abels, Erik R., Sybren L.N. Maas, et al. 2019. “Glioblastoma-Associated Microglia Reprogramming Is Mediated by Functional Transfer of Extracellular MiR-21.” Cell Reports 28(12): 3105–3119.e7. https://doi.org/10.1016/j.celrep.2019.08.036.

Abels, Erik R., Marike L.D. Broekman, Xandra O. Breakefield, and Sybren L.N. Maas. 2019. “Glioma EVs Contribute to Immune Privilege in the Brain.” Trends in Cancer 5(7): 393–96.

Albanese, Manuel et al. 2021. “MicroRNAs Are Minor Constituents of Extracellular Vesicles That Are Rarely Delivered to Target Cells.” PLOS Genetics 17(12): e1009951. https://doi.org/10.1371/journal.pgen.1009951.

Al Ali, Jamal et al. 2021. “TAF1 Transcripts and Neurofilament Light Chain as Biomarkers for X-Linked Dystonia-Parkinsonism.” Movement disorders : official journal of the Movement Disorder Society 36(1): 206–15.

Badhwar, Aman Preet, and Arsalan S. Haqqani. 2020. “Biomarker Potential of Brain-Secreted Extracellular Vesicles in Blood in Alzheimer’s Disease.” Alzheimer’s and Dementia: Diagnosis, Assessment and Disease Monitoring 12(1). /pmc/articles/PMC7085285/ (May 20, 2021).

Balaj, Leonora et al. 2011. “Tumour Microvesicles Contain Retrotransposon Elements and Amplified Oncogene Sequences.” Nature communications 2: 180.

Barman, Bahnisikha et al. 2022. “VAP-A and Its Binding Partner CERT Drive Biogenesis of RNA-Containing Extracellular Vesicles at ER Membrane Contact Sites.” Developmental Cell 57(8): 974–994.e8. https://www.sciencedirect.com/science/article/pii/S1534580722002064.

Benedikter, Birke J et al. 2017. “Ultrafiltration Combined with Size Exclusion Chromatography Efficiently Isolates Extracellular Vesicles from Cell Culture Media for Compositional and Functional Studies.” Scientific reports 7(1): 15297.

Bhaskaran, Vivek et al. 2019. “The Functional Synergism of MicroRNA Clustering Provides Therapeutically Relevant Epigenetic Interference in Glioblastoma.” Nature communications 10(1): 442.

Bolukbasi, Mehmet Fatih et al. 2012. “MiR-1289 and ‘Zipcode’-like Sequence Enrich MRNAs in Microvesicles.” Molecular Therapy - Nucleic Acids 1: e10. https://www.sciencedirect.com/science/article/pii/S216225311630066X.

Breyne, Koen et al. 2022. “Exogenous Loading of Extracellular Vesicles, Virus-like Particles, and Lentiviral Vectors with Supercharged Proteins.” Communications Biology 5(1).

Budnik, Vivian, Catalina Ruiz-Cañada, and Franz Wendler. 2016. “Extracellular Vesicles Round off Communication in the Nervous System.” Nature Reviews Neuroscience 17(3): 160–72. http://www.nature.com/articles/nrn.2015.29.

Carmona, Vitor et al. 2017. “Unravelling Endogenous MicroRNA System Dysfunction as a New Pathophysiological Mechanism in Machado-Joseph Disease.” Molecular Therapy 25(4): 1038–55. https://www.sciencedirect.com/science/article/pii/S1525001617300552.

Chivet, Mathilde et al. 2014. “Exosomes Secreted by Cortical Neurons upon Glutamatergic Synapse Activation Specifically Interact with Neurons.” Journal of extracellular vesicles 3: 24722.

Chu, Aimee J, and Joanna M Williams. 2021. “Astrocytic MicroRNA in Ageing, Inflammation, and Neurodegenerative Disease.” Frontiers in physiology 12: 826697.

Dautzenberg, Iris J.C., Martijn J.W.E. Rabelink, and Rob C. Hoeben. 2021. “The Stability of Envelope-Pseudotyped Lentiviral Vectors.” Gene Therapy 28(1–2): 89–104. http://dx.doi.org/10.1038/s41434-020-00193-y.

Van Duyne, Gregory D. 2015. “Cre Recombinase.” Microbiology spectrum 3(1): MDNA3-0014–2014.

Emmanouilidou, Evangelia et al. 2010. “Cell-Produced Alpha-Synuclein Is Secreted in a Calcium-Dependent Manner by Exosomes and Impacts Neuronal Survival.” The Journal of neuroscience : the official journal of the Society for Neuroscience 30(20): 6838–51.

England, Christopher G, Emily B Ehlerding, and Weibo Cai. 2017. “NanoLuc: A Small Luciferase Is Brightening Up the Field of Bioluminescence.” Physiology & behavior 176(3): 139–48.

Fader, Claudio M, Diego Sánchez, Marcelo Furlán, and María I Colombo. 2008. “Induction of Autophagy Promotes Fusion of Multivesicular Bodies with Autophagic Vacuoles in K562 Cells.” Traffic (Copenhagen, Denmark) 9(2): 230–50.

Frühbeis, Carsten et al. 2013. “Neurotransmitter-Triggered Transfer of Exosomes Mediates Oligodendrocyte-Neuron Communication.” PLoS Biology 11(7).

Garcia-Martin, Ruben et al. 2022. “MicroRNA Sequence Codes for Small Extracellular Vesicle Release and Cellular Retention.” Nature 601(7893): 446–51. https://doi.org/10.1038/s41586-021-04234-3.

Gupta, Dhanu, Antje Maria Zickler, and Samir El Andaloussi. 2021. “Dosing Extracellular Vesicles.” Advanced Drug Delivery Reviews 178: 113961. https://doi.org/10.1016/j.addr.2021.113961.

Hall, Mary P et al. 2012. “Engineered Luciferase Reporter from a Deep Sea Shrimp Utilizing a Novel Imidazopyrazinone Substrate.” ACS Chemical Biology 7(11): 1848–57. https://doi.org/10.1021/cb3002478.

Hill, Andrew F. 2019. “NeuroEVs : Characterizing Extracellular Vesicles Generated in the Neural Domain, by Extracellular Vesicles and Neurodegenerative Diseases.” Journal of Neuroscience 39(47): 9269–73.

Houseley, Jonathan, and David Tollervey. 2009. “The Many Pathways of RNA Degradation.” Cell 136(4): 763–76. https://www.sciencedirect.com/science/article/pii/S0092867409000671.

Huang, Yiyao et al. 2020. “Influence of Species and Processing Parameters on Recovery and Content of Brain Tissue-Derived Extracellular Vesicles.” Journal of Extracellular Vesicles 9(1).

Jeppesen, Dennis K. et al. 2019. “Reassessment of Exosome Composition.” Cell 177(2): 428–445.e18. https://doi.org/10.1016/j.cell.2019.02.029.

Khattar, Khattar E, Janice Safi, Anne-Marie Rodriguez, and Marie-Luce Vignais. 2022. “Intercellular Communication in the Brain through Tunneling Nanotubes.” Cancers 14(5).

Kojima, Ryosuke et al. 2018. “Designer Exosomes Produced by Implanted Cells Intracerebrally Deliver Therapeutic Cargo for Parkinson’s Disease Treatment.” Nature Communications 9(1). http://dx.doi.org/10.1038/s41467-018-03733-8.

Lang, Ming-Fei et al. 2012. “Genome-Wide Profiling Identified a Set of MiRNAs That Are Differentially Expressed in Glioblastoma Stem Cells and Normal Neural Stem Cells.” PloS one 7(4): e36248.

Li, Mu et al. 2014. “Analysis of the RNA Content of the Exosomes Derived from Blood Serum and Urine and Its Potential as Biomarkers.” Philosophical transactions of the Royal Society of London. Series B, Biological sciences 369(1652).

Li, Qingyun, and Ben A Barres. 2018. “Microglia and Macrophages in Brain Homeostasis and Disease.” Nature reviews. Immunology 18(4): 225–42.

Liu, Haisheng et al. 2022. “Analysis of Extracellular Vesicle DNA at the Single-Vesicle Level by Nano-Flow Cytometry.” Journal of extracellular vesicles 11(4): e12206.

Madisen, Linda et al. 2010. “A Robust and High-Throughput Cre Reporting and Characterization System for the Whole Mouse Brain.” Nature Neuroscience 13(1): 133–40. https://doi.org/10.1038/nn.2467.

Mahjoum, Shadi, David Rufino-ramos, and Marike L D Broekman. 2021. “Living Proof of Activity of Extracellular Vesicles in the Central Nervous System.”

Men, Yuqin et al. 2019. “Exosome Reporter Mice Reveal the Involvement of Exosomes in Mediating Neuron to Astroglia Communication in the CNS.” Nature Communications 10(1).

Morel, Lydie et al. 2013. “Neuronal Exosomal MiRNA-Dependent Translational Regulation of Astroglial Glutamate Transporter GLT1 * □.” Journal of Biological Chemistry 288(10): 7105– 16. http://dx.doi.org/10.1074/jbc.M112.410944.

van Niel, Guillaume, Gisela D’Angelo, and Graça Raposo. 2018. “Shedding Light on the Cell Biology of Extracellular Vesicles.” Nature Reviews Molecular Cell Biology 19(4): 213–28. https://doi.org/10.1038/nrm.2017.125.

Nolte-’t Hoen, Esther N M, et al. 2012. “Deep Sequencing of RNA from Immune Cell-Derived Vesicles Uncovers the Selective Incorporation of Small Non-Coding RNA Biotypes with Potential Regulatory Functions.” Nucleic acids research 40(18): 9272–85.

Norman, Maia et al. 2021. “L1CAM Is Not Associated with Extracellular Vesicles in Human Cerebrospinal Fluid or Plasma.” Nature Methods 18(6): 631–34. http://dx.doi.org/10.1038/s41592-021-01174-8.

Norrman, Karin et al. 2010. “Quantitative Comparison of Constitutive Promoters in Human ES Cells.” PloS one 5(8): e12413.

O’Brien, Killian et al. 2020. “RNA Delivery by Extracellular Vesicles in Mammalian Cells and Its Applications.” Nature Reviews Molecular Cell Biology 21(10): 585–606. https://doi.org/10.1038/s41580-020-0251-y.

Parr-Brownlie, Louise C. et al. 2015. “Lentiviral Vectors as Tools to Understand Central Nervous System Biology in Mammalian Model Organisms.” Frontiers in Molecular Neuroscience 8(MAY): 1–12.

Patel, B. et al. 2016. “Exosomes Mediate the Acquisition of the Disease Phenotypes by Cells with Normal Genome in Tuberous Sclerosis Complex.” Oncogene 35(23): 3027–36.

Pegtel, D Michiel, et al. 2010. “Functional Delivery of Viral MiRNAs via Exosomes.” Proceedings of the National Academy of Sciences of the United States of America 107(14): 6328–33.

Rajendran, Lawrence et al. 2006. “Alzheimer’s Disease Beta-Amyloid Peptides Are Released in Association with Exosomes.” Proceedings of the National Academy of Sciences of the United States of America 103(30): 11172–77.

Ridder, Kirsten et al. 2014. “Extracellular Vesicle-Mediated Transfer of Genetic Information between the Hematopoietic System and the Brain in Response to Inflammation.” 12(6).

Ridder, Kirsten et al. 2015. “Extracellular Vesicle-Mediated Transfer of Functional RNA in the Tumor Microenvironment.” OncoImmunology 4(6): 1–8.

Rufino-ramos, David et al. 2022. “Biomaterials Using Genetically Modified Extracellular Vesicles as a Non-Invasive Strategy to Evaluate Brain-Specific Cargo.” Biomaterials 281: 121366. https://doi.org/10.1016/j.biomaterials.2022.121366.

Ruivo, Carolina F et al. 2022. “Extracellular Vesicles from Pancreatic Cancer Stem Cells Lead an Intratumor Communication Network (EVNet) to Fuel Tumour Progression.” Gut: gutjnl-2021–324994.

Rustom, Amin et al. 2004. “Nanotubular Highways for Intercellular Organelle Transport.” Science (New York, N.Y.) 303(5660): 1007–10.

Shi, Min et al. 2019. “New Windows into the Brain: Central Nervous System-Derived Extracellular Vesicles in Blood.” Progress in Neurobiology 175(July 2018): 96–106. https://doi.org/10.1016/j.pneurobio.2019.01.005.

Shurtleff, Matthew J et al. 2016. “Y-Box Protein 1 Is Required to Sort MicroRNAs into Exosomes in Cells and in a Cell-Free Reaction” ed. Timothy W Nilsen. eLife 5: e19276. https://doi.org/10.7554/eLife.19276.

Steenbeek, Sander C et al. 2018. “Cancer Cells Copy Migratory Behavior and Exchange Signaling Networks via Extracellular Vesicles.” The EMBO Journal 37(15): 1–20.

Sterzenbach, Ulrich et al. 2017. “Engineered Exosomes as Vehicles for Biologically Active Proteins.” Molecular Therapy.

Street, Jonathan M et al. 2012. “Identification and Proteomic Profiling of Exosomes in Human Cerebrospinal Fluid.” Journal of Translational Medicine 10(1): 5. https://doi.org/10.1186/1479-5876-10-5.

Su, Huaqi et al. 2021. “Characterization of Brain-Derived Extracellular Vesicle Lipids in Alzheimer’s Disease.” Journal of Extracellular Vesicles 10(7).

Ter-Ovanesyan, Dmitry et al. 2021. “Framework for Rapid Comparison of Extracellular Vesicle Isolation Methods.” eLife 10: 1–17.

Théry, Clotilde et al. 2018. “Minimal Information for Studies of Extracellular Vesicles 2018 (MISEV2018): A Position Statement of the International Society for Extracellular Vesicles and Update of the MISEV2014 Guidelines.” Journal of Extracellular Vesicles 7(1).

Tóth, Eszter et al. 2021. “Formation of a Protein Corona on the Surface of Extracellular Vesicles in Blood Plasma.” Journal of Extracellular Vesicles 10(11).

Valadi, Hadi et al. 2007. “Exosome-Mediated Transfer of MRNAs and MicroRNAs Is a Novel Mechanism of Genetic Exchange between Cells.” Nature cell biology 9(6): 654–59.

Vassileff, Natasha et al. 2020. “Revealing the Proteome of Motor Cortex Derived Extracellular Vesicles Isolated from Amyotrophic Lateral Sclerosis Human Postmortem Tissues.” Cells 9(7).

Vella, Laura J. et al. 2017. “A Rigorous Method to Enrich for Exosomes from Brain Tissue.” Journal of Extracellular Vesicles 6(1). https://doi.org/10.1080/20013078.2017.1348885.

Villarroya-Beltri, Carolina et al. 2013. “Sumoylated HnRNPA2B1 Controls the Sorting of MiRNAs into Exosomes through Binding to Specific Motifs.” Nature Communications 4(1): 2980. https://doi.org/10.1038/ncomms3980.

Van Der Vos, Kristan E., et al. 2016. “Directly Visualized Glioblastoma-Derived Extracellular Vesicles Transfer RNA to Microglia/Macrophages in the Brain.” Neuro-Oncology 18(1): 58–69.

Wang, Yipeng et al. 2017. “The Release and Trans-Synaptic Transmission of Tau via Exosomes.” Molecular neurodegeneration 12(1): 5.

Wei, Zhiyun et al. 2017. “Coding and Noncoding Landscape of Extracellular RNA Released by Human Glioma Stem Cells.” Nature Communications 8(1): 1145. https://doi.org/10.1038/s41467-017-01196-x.

Wettergren, Erika Elgstrand, Fredrik Gussing, Luis Quintino, and Cecilia Lundberg. 2012. “Novel Disease-Specific Promoters for Use in Gene Therapy for Parkinson’s Disease.” Neuroscience Letters 530(1): 29–34. https://www.sciencedirect.com/science/article/pii/S0304394012013213.

You, Yang et al. 2022. “Human Neural Cell Type-specific Extracellular Vesicle Proteome Defines Disease-related Molecules Associated with Activated Astrocytes in Alzheimer’s Disease Brain.” Journal of Extracellular Vesicles 11(1).

Zappulli, Valentina et al. 2016. “Extracellular Vesicles and Intercellular Communication within the Nervous System.” Journal of Clinical Investigation 126(4): 1198–1207.

Zomer, Anoek et al. 2015. “In Vivo Imaging Reveals Extracellular Vesicle-Mediated Phenocopying of Metastatic Behavior.” Cell 161(5): 1046–57.

Zomer, Anoek, Sander Christiaan Steenbeek, Carrie Maynard, and Jacco Van Rheenen. 2016. “Studying Extracellular Vesicle Transfer by a Cre-LoxP Method.” Nature Protocols 11(1): 87– 101.

